# SIRPα+ PD-L1+ bone marrow macrophages aid AML growth by modulating T cell function

**DOI:** 10.1101/2024.09.15.613123

**Authors:** Flora Birch, Sara Gonzalez-Anton, Myriam L.R. Haltalli, Probir Chakravarty, Federica Bruno, Christiana Georgiou, Cera Mai, Yvette Todd, Beth Payne, Sabine Blum, Constandina Pospori, Caroline Arber, Richard Burt, Floriane S. Tissot, Cristina Lo Celso

## Abstract

Acute myeloid leukemia (AML) continues to have a poor prognosis due to its ability to relapse following initial response to chemotherapy. While immunotherapies hold the promise to revolutionize cancer treatment, AML has been particularly challenging to target. It is therefore important to better understand the relationship between AML cells and immune cells within the bone marrow (BM) microenvironment, where this disease grows. Here we focus on non-malignant BM macrophages, and using a combination of intravital microscopy, flow cytometry, transcriptomics and functional analyses we identify a subpopulation of immunomodulatory BM macrophages (IMMs) with a unique profile and function during AML progression. While the majority of macrophages are already being lost at early infiltration, IMMs are locally enriched. They are capable of efferocytosis and support AML growth through inhibition of T cells. Enrichment of IMMs in the BM of patients developing early relapse indicates that future development of interventions that target IMMs’ development and function may improve AML patients’ outcome.

## Introduction

Acute myeloid leukemia (AML) remains one of the deadliest forms of cancer. Chemotherapy continues to be the first line treatment of choice, compared to other cancers where targeted drugs and immunotherapy approaches are more established in upfront treatment. Despite the recent development of a few targeted therapeutic approaches, refractory disease or relapse after therapy is frequent (50-90% depending on subtypes) and is often resistant to available treatments(1,2). An important hallmark of AML is its capacity to evade immune surveillance(3,4). A number of cell-intrinsic mechanisms have been described to be involved in immune evasion. These include the loss of human leukocyte antigens (HLA) molecules, which are essential for antigen presentation(3,5,6); expression of inhibitory ligands that hinder T-cell responses(7); and production of inhibitory ligands that impede macrophage-mediated phagocytosis(8). While these leukemia-intrinsic mechanisms all contribute towards an immunosuppressive response, extrinsic mechanisms are also believed to play a role in disease progression, but remain poorly understood. Therefore, it is essential to identify the cells responsible for creating such an immunosuppressive environment. A better understanding of the AML bone marrow (BM) microenvironment can pave the way for the development of more targeted therapies with fewer side effects and improved efficacy in preventing relapse.

Within the BM environment, it is clear that macrophages play a crucial role in AML development as their abundance, measured in single cell transcriptomics datasets, is associated with a poor outcome(9); however, the mechanisms underpinning this phenomenon are not clear. It is well accepted that macrophages are heterogeneous, and the initial classificaiton of M1 and M2 (pro and anti-inflammatory) subtypes has been surpassed as multiple sub-populations and activation states of macrophages have been described in several tissues(10). However, there is a lack of consensus regarding the classification of BM macrophages in the context of AML in both mice and humans. CD11b+ Ly6G-MHCII-Ly6C-macrophages were shown to be elevated in the BM of leukaemic mice compared to non-leukaemic mice(11). Similarly, GR1^lo^ MCSFR^int^ F4/80^hi^ SSC^lo^ macrophages isolated from the BM of AML burdened mice were found to be enriched for M1 gene signatures(12). In contrast, CD11b-F4/80+ CD169+ VCAM1+ macrophages were shown to display a reduction in M1 markers, including MHC class II and CD80(13). While these studies have focused on understanding the direct impact of AML on BM macrophages and correlate it to AML outcome, the function of the different macrophage populations in the context of AML remains unknown. The recent disappointing results of clinical trials with the macrophage-targeted immune checkpoint inhibitor Magrolimab(14) highlight both the challenge and importance of understanding the interplay between macrophage subpopulations and AML cells.

Here, we identify a sub-population of macrophages already present in healthy BM and with a unique function in AML progression. These cells, named immunomodulatory macrophages (IMMs), express SIRPα and PD-L1 and locally increase within AML cell clusters at early stages of AML growth, while all other macrophage subsets, collectively named hematopoiesis supportive macrophages (HSMs), are rapidly lost. Combining different mouse models and using clodronate liposomes to deplete macrophages, we show that targeting IMMs specifically delays AML growth in a T cell dependent manner. Single cell RNA-sequencing (scRNA-seq) analysis combined with intravital microscopy (IVM) and *in vitro* co-cultures revealed IMMs to be efferocytic cells capable of uptaking AML cells and inhibiting T cells. In the murine model, these macrophages were the only ones to express CD206, a marker identified in patients to be associated with poor survival(15). Analysis of paired human AML BM samples collected at diagnosis and post-chemotherapy indicated two populations of macrophages similar to those identified in mice. Patients with a higher fraction of immunosuppressive macrophages post-chemotherapy went on to develop earlier relapse, highlighting the importance of this macrophage population in disease progression.

## Methods

### Animal Studies

All animal work was carried out in accordance with the animal ethics committee (Animal Welfare Ethical Review Body) at Imperial College London and Sir Francis Crick institute and UK Home Office regulations (Animals (Scientific Procedures) Act, 1986) under license numbers 70/8403 and PP9504146.

CD169-DTR and CD169-iCre mice originated from Masato Tanaka (RIKEN Research Center for Allergy and Immunology) were imported and bred at the Sir Francis Crick Institute. CD169-Cre mice were crossed with Rosa26-eYFP mice donated by Frank Costantini (University of Columbia) and bred by the Francis Crick Institute. The TCRαKO and mT/mG lines were imported from Jackson Laboratories and bred by the Francis Crick Institute. C57BL/6 mice were obtained from Charles River. Female mice >8 weeks of age were used in all of the experiments. The mice were housed in Tecniplast mouse greenline cages with appropriate bedding and enrichment. The temperature, humidity and light cycles were kept within the UK Home Office code of practice, with the temperature between 20 and 24 °C, the room humidity at 45–65% and a 12 h/12 h light/dark cycle with a 30-min dawn and dusk period to provide a gradual change.

### Administration of drugs, liposomes and macrophage depleting agents

For Diphtheria Toxin (DT) treatment, DT (Sigma-Aldrich) was injected intraperitoneally (10 ng/g body weight) for 2 consecutive days followed by injections every 3 days for up to three weeks into CD169-DTR mice or WT littermates (control).

For clodronate treatment, clodronate liposomes or PBS liposomes (Liposoma BV) were injected intravenously (i.v., 0.1 ml/10 g body weight) every 4 days for up to 18 days.

For chemotherapy, 100 mg of Cytarabine (MedChemExpress) was dissolved in 10% Dimethyl sulfoxide (DMSO), 40% PEG300, 5% Tween-80 and 45% PBS to achieve a stock concentration of 100 mg/ml. 100 mg of Doxorubicine (MedChemExpress) was dissolved in 5% DMSO, 40% PEG-300, 5% Tween-80 and 50% PBS to achieve a stock concentration of 2 mg/ml. When AML infiltration was greater than 10% in the blood (an indication of BM infiltration >60%) (18), as measured by flow cytometry, 100 mg/kg body weight of Cytarabine and 3mg/kg body weight of Doxorubicine were injected i.v. for 3 days followed by 2 days of injections with only Cytarabine, consistent with the commonly used 5+3 AML drug regimen (16).

### Leukemia transplantation experiments

Murine AML cells were generated as described in Duarte et al. (16). Briefly, granulocyte macrophage progenitors (GMPs) were sort purified from mT/mG transgenic mice, transduced with pMSCV-MLL-AF9-GFP retroviruses as described by Krivtsov and colleagues (17) and transplanted into sub lethally irradiated mice (two doses of 3.3 Gy, at least 3 hours apart). At 8+ weeks post-transplantation, recipient mice developed leukemia characterized by multi-organ infiltration of Tomato+ cells. Tomato+ cells were harvested from BM and spleen and cells from each primary recipient were labelled as a separate batch and cryopreserved. Primary cells from different batches were thawed, suspended in PBS and 100,000 viable cells were injected i.v. into secondary, non-conditioned recipient mice. In some experiments, cells from secondary, untreated recipients were used. Progressive AML expansion was observed from day 8-10 and full BM infiltration was typically reached around day 20-24, with some variability depending on the batch and passage used (16,18). This was accompanied by infiltration of the spleen and blood, typically delayed compared to BM infiltration.

### Flow cytometry analysis and sorting

For macrophage analysis of murine BM samples, bones were crushed in PBS with 2% fetal bovine serum (FBS) and the cells were filtered through a 70 µm strainer, depleted of red blood cells (18) and stained with relevant antibodies. For macrophage analysis of human BM samples, BM aspirate samples were thawed in 100% FBS, spun and resuspended in 2% FBS and stained with relevant antibodies. For information on all of the antibodies used, see Supplementary Table S1 and S2. For analysis of AML infiltration in murine blood, 50 or 500 µl of blood was obtained by tail vein venipuncture or cardiac puncture and mixed with ethylenediaminetetraacetic acid (EDTA) to prevent clotting. Red blood cells were subsequently lysed and the cells were washed with 2% FBS. Cells were analyzed with an LSRFortessa (BD Biosciences) or Cytek Aurora (Cytek) or sorted on a FACSAriaIII (BD Biosciences) and the data were analyzed with FlowJo and the FlowSOM, t-SNE and UMAP plugins (Tree Star). Supplementary Figures 2 & 8 contain the gating strategies used for analysis of the data presented in this manuscript.

### AML cell culture and induction of apoptosis

Primary mT/mG leukemia cells were thawed and plated at 5×10^5^ cells/well in a 6-well tissue culture non-treated plate in Roswell Park Memorial Institute (RPMI) 1640 medium, 10% FBS, 2 mM L-Glutamine (Thermo Fisher), 10 ng/ml human interleukin-6 (hIL-6) (Peprotech), 10 ng/mL murine stem cell factor (m-SCF) (Peprotech) and 6 ng/mL murine interleukin-3 (mIL-3) (Peprotech). Every two days, cells were counted using a hematocytometer and plated at 1×10^5^ cells/ml in fresh medium. Apoptosis induction was performed by treating AML cells with 0.01µg/mL Doxorubicin (MedChemExpress) for 24 hours. Apoptosis was confirmed by flow cytometry with >60% of cells being Annexin V+ (BD Biosciences), used following the manufacturer’s reagent instructions.

### Differentiation of sort purified monocytes into IMMs

Following BM harvest from a healthy mouse and staining as described above, 1×10^6^ CD11b+ Ly6G-Ly6C^hi^ cells were sort purified at 4^◦^C in PBS with 10% FBS. The cells were centrifuged at 300 x g for 10 minutes, resuspended in macrophage media composed of Dulbecco’s Modified Eagle Medium/Nutrient Mixture F-12 (DMEM F12) supplemented with 10% FBS, 50 ng/mL of M-CSF (Peprotech), 10mM of L-Glutamine (ThermoFisher) and 1% Penicillin/Streptomycin (Life Technologies) and plated at 2×10^5^/mL in a flat bottom 96 well-plate. The plates were incubated at 37^◦^C for 48 hours. Following incubation, the cells were harvested using a cell scraper (Greiner Bio-One Ltd) in 200 μl of fresh PBS. The cells were centrifuged and stained for flow cytometry analysis and IMMs phenotype was confirmed through the expression of F4/80, SIRPα and PD-L1 and reduced expression of Ly6C.

### AML uptake assay

mT/mG AML cells were labelled with 1 μl/ml of cell trace violet (CTV) (ThermoFisher) and resuspended at 2×10^5^/mL. Co-cultures of *in vitro* differentiated IMMs and HSMs with live and apoptotic CTV+ AML cells were carried out at 3 different ratios, 2:1, 1:1 and 1:2 macrophage:AML cells in macrophage medium for 24 hours. CTV levels in macrophages were assessed by flow cytometry and were the readout for AML uptake.

### Macrophages and T cells co-cultures

On the day prior to the co-culture, flat bottom 96 well plates were coated with 50 μL of 1% anti-CD3 antibody (BD Biosciences) diluted in sterile PBS. On the day of the experiment, splenic T cells were isolated using EasySep Mouse T cell isolation kit (StemCell, Cat. No. 19851) and labelled with CTV (Invitrogen) according to the manufacturers’ instructions. Co-culture of *in vitro* differentiated IMMs or HSMs with T cells were setup at a ratio of 0.4:1, 0.2:1, 0.1:1, 0.05:1 and 0.025:1 macrophages:T cells in macrophage medium for 72 hours. To activate the T cells, 1 µg/ml of anti-CD28 antibody (BD Biosciences) was added. As a positive control, T cells were incubated alone with anti-CD3/CD28 antibodies. As a negative control, T cells were incubated alone without antibodies.

T cell proliferation and division index was analyzed by flow cytometry using the FlowJo Proliferation Tool and T cell absolute number was calculated using Calibrite beads (BD Bioscience).

### Intravital microscopy (IVM)

IVM was performed using Zeiss LSM 780 and 980 upright confocal and two photon hybrid microscopes equipped with 450, 488, 514, 561 and 633nm excitation lasers, a tunable multiphoton laser (Spectraphysics Mai Tai eDeep See, 680-1020nm and Insight dual line 1040nm fixed and 680-1300 tunable, respectively), four non-descanned detectors (NDD) and an internal detector array. Signal was visualized using a Zeiss W Plan-Apochromat 20X DIC water immersion lens (1.0 NA). Live imaging of the calvarium BM was carried out as described in previously published reports(16,20). CD169-YFP fluorescent protein was excited with the 488 nm laser while Tomato protein was excited with the 561 nm laser. SIRPα expression was visualized by injecting 25 µg of anti-SIRPα APC conjugated antibody and excited with the 633 nm laser.

### Single cell RNA sequencing

For the 10× Chromium (10× Genomics) experiments, two mice were selected based on AML infiltration to be representative of an early timepoint (AML infiltration between 10-20% in the BM). Tomato-CD11b+ Ly6G-Ly6C^lo^ SSCA^lo^ F4/80+ cells were sort purified, as described above, from two AML burdened mice and two control mice. The cells from each condition were processed according to the manufacturer’s protocol. Library generation for 10x Chromium analysis was performed following the Chromium Single Cell 3′ Reagents Kits (10x Genomics) and sequenced on an Hiseq4000 (Illumina), to achieve an average of ∼44,000 reads per cell and ∼11,000 cells per sample. Raw reads were initially processed by the Cell Ranger v.6.1.2 pipeline (21), which deconvolved reads to their cell of origin using the UMI tags, aligned these to the mm10 transcriptome using STAR (v.2.5.1b) (22) and reported cell-specific gene expression count estimates. All subsequent analyses were performed in R v.4.1.0 (23) using the Seurat (v4) package (24). Genes were considered to be ‘expressed’ if the estimated (log10) count was at least 0.1. Primary filtering was then performed as follows: a sample-specific threshold for low-quality cells was identified using median absolute deviation (MAD) measures for cells expressing >3 MAD of mitochondrial gene expression, which were removed from further analyses; cells expressing fewer than 500 genes were also removed. Four samples (2 Ctrl_Macrophages and 2 AML_Macrophages) were integrated using Seurat’s ‘IntegrateData’ function, after identifying 2000 anchor features. PCA decomposition was performed using 50PCs based on a JackStraw determined signficant of P < 0.01.

UMAP analysis was performed on the intergated data at a resolution of 0.2, which produced 16 clusters. Clusters were annotated using the scMCA package [https://github.com/ggjlab/scMCA] using data from the single cell MouseCellAtlas [https://bis.zju.edu.cn/MCA/]. Macrophage clusters were identified and re-clustered. For data visualisation, dimensional reduction of the integrated Macrophage-clusters-only dataset was achieved using UMAP with 30 principal components, with a resolution of 0.2 producing 6 clusters (Fig. 3B). For each gene, expression levels are normalized by the total expression, multiplied by a scale factor (10,000) and log-transformed. Cluster markers were identified using FindAllMarkers function in Seurat (Wilcoxon rank sum test). Differential gene expression between clusters or the same cluster comparing samples were identified using FindMarkers function in Seurat (Wilcoxon rank sum test). Differentially expressed genes were ranked by their log2FC value and used for Gene Set Enrichment Analysis using hallmark, pathways and processes gene sets from MsigDB (v6)(25) or tested for overenrichment analysis using the clusterProfiler package(26).

### Human Samples

BM aspirate samples from the UCLH biobank (REC reference 20/YH/0088) or from the CHUV (CER-VD project number: 2018-01951) were obtained following patient’s consent, mononuclear cells isolated by density gradient centrifugation (Ficoll; Amersham Biosciences, Little Chalfont, United Kingdom) and stored in liquid nitrogen according to Human Tissue Act guidelines. Samples were thawed and washed with 100% FBS, then 50% FBS:50% RPMI prior to flow cytometry on the same day. Samples were collected at diagnosis and/or following peripheral blood count recovery after the first cycle of induction chemotherapy. Supplementary Table S3 provides further, anonymised, information on the samples and the chemotherapy protocols used.

### Statistical analyses

Raw data were analyzed using Excel (Microsoft) and GraphPad Prism (GraphPad Software Inc.). t-tests were performed to compare group means when a single comparison was needed. In the case where statistical analysis of multiple comparisons was needed, one-way ANOVA test was used. All differences were considered statistically significant when p<0.05. ∗ represents p<0.05, ∗∗ represents p<0.01, ∗∗∗ represents p<0.001 and ∗∗∗∗ represents p<0.0001. Detailed statistical analysis and the number of biological and technical replicates for each experiment can be found in the figure legends.

### Data Availability

Sequencing data will be made publicly available upon publication.

## Results

### Macrophages are enriched inside AML cell clusters at early disease stages

To explore the dynamics of BM macrophages during AML growth we took advantage of the well-established murine model of AML driven by overexpression of the MLL-AF9 oncogenic translocation and capable to grow in non-conditioned, immunocompetent C57Bl/6 syngenic mice (16,18,27). This model is therefore unique in allowing the study of interactions between AML and healthy immune cells in vivo in physiological conditions, something that is impossible to achieve with othr murine or patient-derived xenograft (PDX)-based models. In this model, AML cells are known to grow through progressive expansion of localized foci, which can be exemplified through a few stages (Fig. 1A). Following intravenous injection of 100,000 AML cells, the engrafting stage corresponds to an overall BM infiltration of 0-5% measured by flow cytometry, where mostly single AML cells are detectable by microscopy. During the early (BM infiltration of 5-20%) and intermediate (BM infiltration of 20-60%) stages, AML cells grow through the expansion of foci that locally remodel the BM microenvironment (16). To explore whether similar localized effects would perturb BM macrophages, we used intravital microscopy and directly observed AML cells and BM macrophages simultaneously in calvarium BM of anesthetized mice. mT/mG AML cells, expressing membrane-bound Tomato (Tomato) fluorescent protein were injected into non-conditioned CD169-Cre x R26-YFP double transgenic mice (shortened to CD169-YFP from here on), a well-established reporter of macrophages (28–31) including in the BM. As a result, both cell types were easily recognizable using confocal microscopy as Tomato and YFP positive, respectively (Fig. 1B).

**Figure 1:**
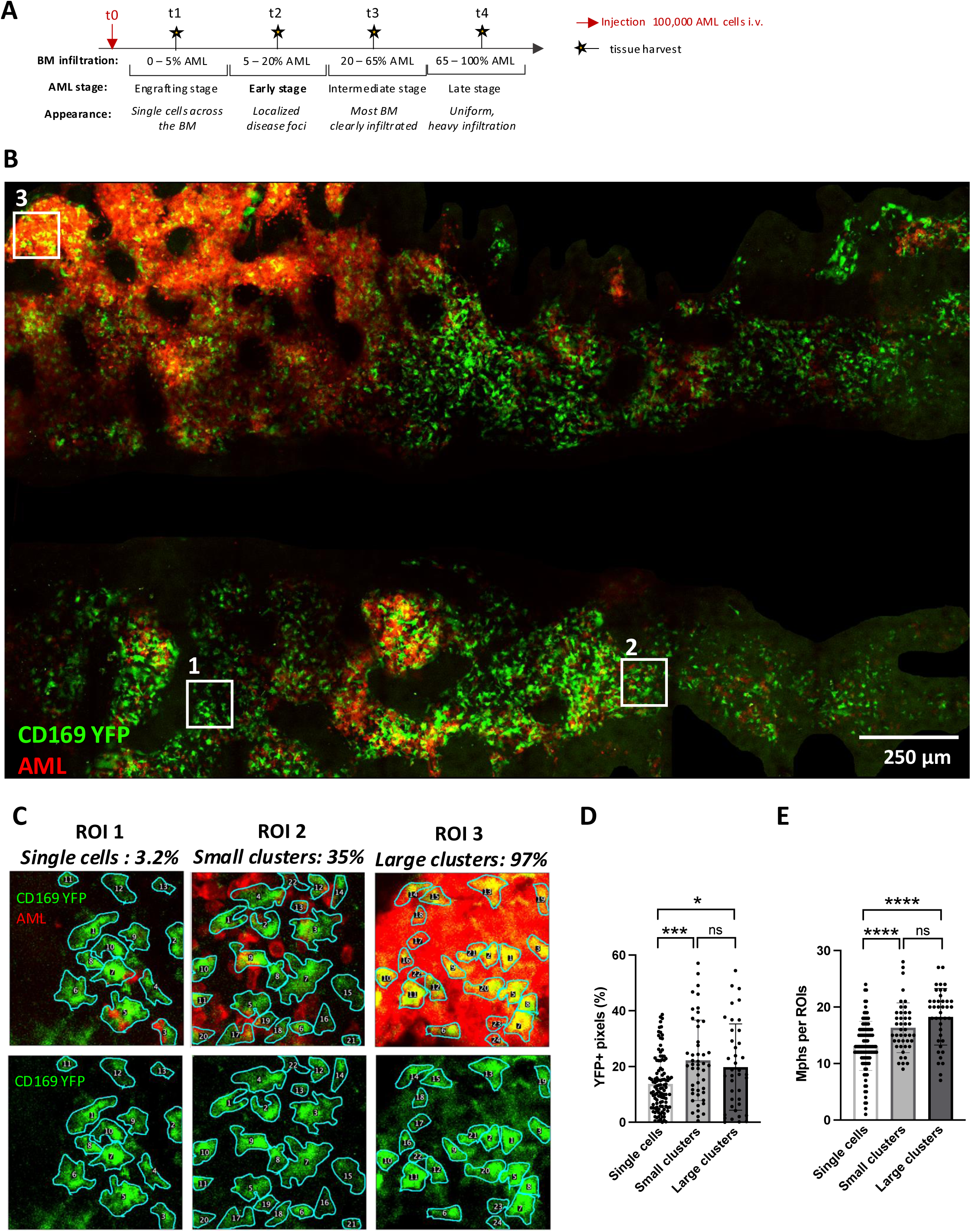
Macrophages are increased inside AML clusters in the bone marrow. **(A)** Schematic detailing AML growth stages and corresponding percentage infiltration in the BM over time. **(B)** Representative calvarium tilescan of early stage AML in CD169-YFP mice. **(C)** Representative ROIs classified based on local infiltration as ‘single cells’ (< 15% Tomato+ pixels pixels), ‘small clusters’ (15 – 50% Tomato+ pixels) and ‘large clusters’ (>50% Tomato+ pixels). The macrophage manual segmentation is outlined in cyan. **(D)** Percentage of YFP+ pixels in each ROI classified according to the fraction of AML in each ROI. **(E)** Number of YFP+ macrophages (Mphs) per ROI classified according to the fraction of AML in each ROI. The statistical significance was determined by one-way ANOVA (n = 3 mice, 112 single cells ROIs, 48 small clusters ROIs and 40 large clusters ROIs = 40). The data are presented as means ±S.D. \**P* < 0.05; \*\*\**P* < 0.0005; \*\*\*\**P* < 0.0001. Scale bar, 250 µm.

Importantly, at early stages of infiltration it is possible to compare the macrophage dynamics in unaffected, lightly and heavily infiltrated BM regions within the same mouse (Fig. 1B and C). Tilescans of calvaria from CD169-YFP mice with <20% AML infiltration showed that YFP+ macrophages were enriched in AML cell clusters compared to non-infiltrated BM regions (Fig. 1B and C). To quantify this enrichment in macrophages, tilescans were divided each into 120µm wide squares, or regions of interest (ROIs), that were classified as: ‘single cells’, ‘small clusters’ and ‘large clusters’ based on the proportion of Tomato+ pixels - a proxy for the number of AML cells-within the ROI (Fig. 1C). As a result, the areas labelled as ‘small clusters’ had low local infiltration, and areas labelled as ‘large clusters’ had high local infiltration. The proportion of YFP+ pixels, corresponding to CD169-YFP macrophages, was significantly increased in ROIs containing AML cell clusters (Fig. 1D). To address whether the increase in YFP+ pixels resulted from an increase in the number of macrophages or an increase in their size, each YFP+ cell was manually segmented. The number of macrophages was significantly higher in ROIs containing AML clusters compared to ROIs containing only sparse AML cells (Fig. 1E). Of note, YFP+ cells were increasingly smaller and more circular (Supplementary Fig. S1A-C). Together, these data show an enrichment in smaller, more rounded macrophages in AML clusters compared to healthy BM areas and raise the question whether AML may be driving these changes in macrophages, and whether all, or a subset, of BM macrophages may be playing a role in disease progression.

### Identification of a BM macrophage sub-population differentially affected by AML growth

To explore whether the increase in macrophages associated with AML clusters corresponded to the enrichment of a subset of macrophages potentially involved in AML progression, the immunophenotype of BM macrophages was characterized using spectral flow cytometry and a high dimensional phenotyping approach (Fig. 2A and B). A broad, 21-colour myeloid marker panel (Fig. 2B and Supplementary Fig. S2) was applied to BM samples harvested from mice at an early stage of AML infiltration (t2, Fig. 1) and healthy controls. An exclusion gating strategy was adopted to remove immune cells other than macrophages, therefore non-myeloid cells (CD11b-), neutrophils (Ly6G^high^), inflammatory monocytes (Ly6C^high^) and eosinophils (SSC-A^high^) were excluded from further analyses (Supplementary Fig. S2A). Among the remaining CD11b+ Ly6G^low/-^ Ly6C^low/-^ SSC-A^low^ cells, macrophages were identified based on their expression of F4/80 (Supplementary Fig. S2A), pooled from all samples and analyzed using a dimensionality reduction approach (Uniform Manifold Approximation and Projection [UMAP] (32)). Among the analyzed cells, further subpopulations clustered separately (Fig. 2C). In particular, one population expressed high levels of PD-L1 and SIRPα and variable levels of TNFα (Fig. 2B and C). All three of these markers are associated with the inflammatory capacity of macrophages (33). We therefore named these cells immunomodulatory macrophages (IMMs) and generated a simplified flow cytometry panel whereby IMMs could be identified based on expression of SIRPα and PD-L1 within the CD11b+, Ly6C^low/-^, F4/80+ myeloid population (Fig. 2D). We broadly named the remaining cells, negative for SIRPα and PD-L1, as hematopoiesis-supportive macrophages (HSMs), as BM macrophages are known to have important roles in HSC retention, iron recycling and erythropoiesis (34–38). Interestingly, while the majority of HSMs were positive for CD169, this marker was expressed more heterogeneously by IMMs, with a skew towards being absent or lowly expressed (Supplementary Fig. S2C).

**Figure 2:**
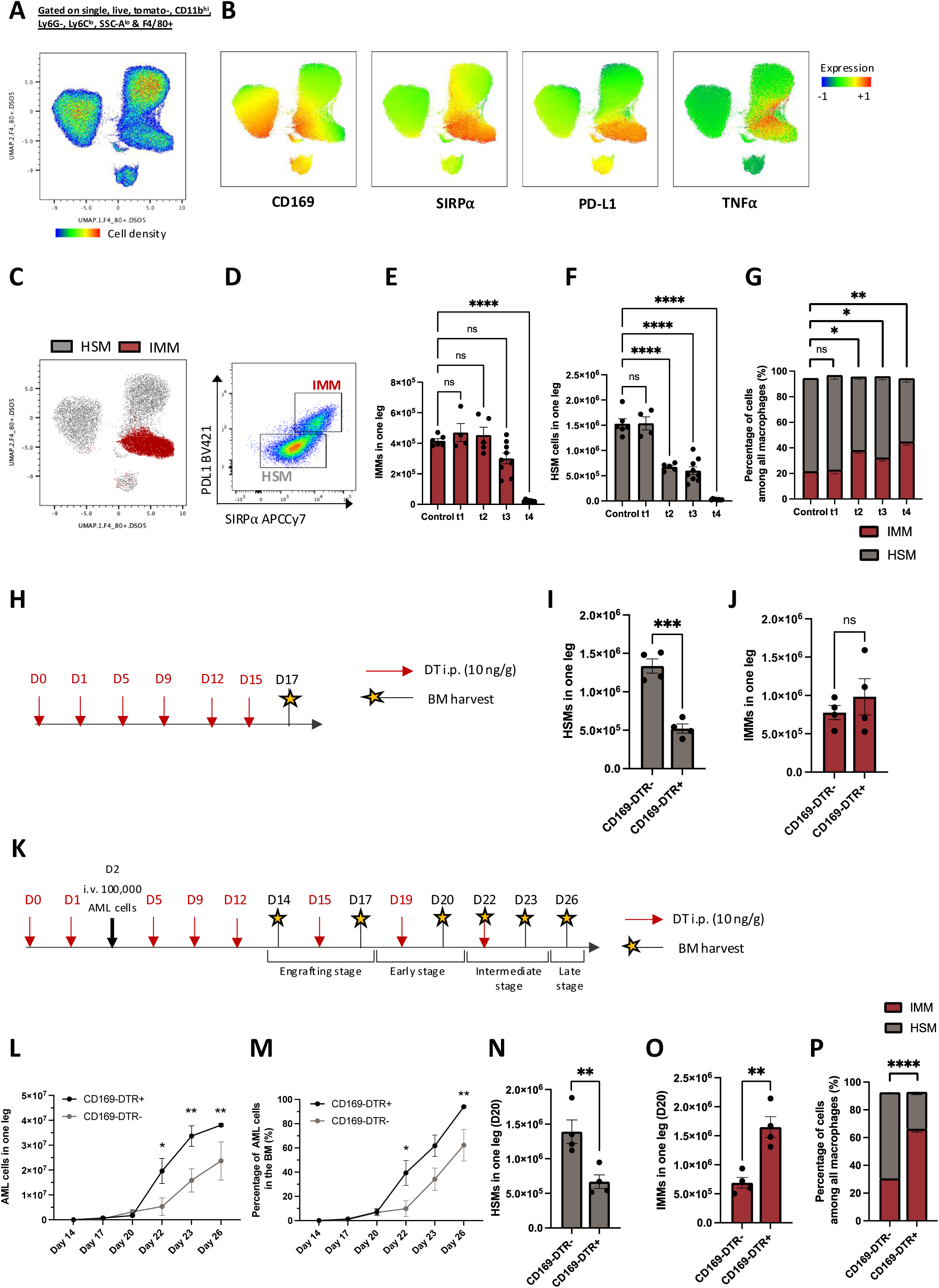
Identification of a sub-population of immunomodulatory macrophages: IMMs. **(A)** Uniform Manifold Approximation and Projection (UMAP) of all single, live, Tomato-, CD11b^hi^, Ly6G-, Ly6C^low^, SSC-A^low^ & F4/80+ cells from control and early AML BM (n= 3 controls and 3 early AML mice). **(B)** Expression of CD169, SIRP⍺, PD-L1 and TNF⍺ in macrophages. **(C)** Localisation of IMMs and HSMs in the UMAP. **(D)** Manual gating strategy for isolating IMMs and HSMs based on PD-L1 and SIRP⍺ expression within F4/80+ cells. **(E-F)** Absolute number of IMMs **(F)** and HSMs **(G)** per leg of mice as AML progresses. **(G)** Proportion of IMMs and HSMs in mice shown in (E). **(H)** Schematic of the DT treatment of CD169-DTR+ and littermate DTR-control mice. **(I-J)** Absolute number of HSMs **(I)** and IMMs **(J)** in DT treated control (n = 4) and CD169-DTR+ (n = 4) mice at day 17 post first injection. **(K)** Schematic of the DT treatment of AML-burdened CD169-DTR+ and littermate control mice. **(L-M)** Absolute number **(L)** and percentage **(M)** of AML cells in the BM. (n = 3-10 mice per group per timepoint). **(N-O)** Absolute number of HSMs **(N)** and IMMs **(O)** in the BM at day 20 post first DT injection. **(P)** Proportion of IMMs and HSMs among all macrophages in the BM at day 20 post first DT injection. The data are presented as means ± s.e.m. \**P* < 0.05; \*\**P* < 0.005; \*\*\**P* < 0.0005; \*\*\*\**P* < 0.0001. The *P* values were determined by two-way ANOVA with post-hoc Bonferroni correction (E-G, L-M) or unpaired two-tailed Student’s *t*-test (I,J, N-P).

To gain some insights into how IMMs and HSMs may be affected by AML growth, we investigated changes in HSMs and IMMs populations size and relative abundance throughout AML progression using flow cytometry of combined femur and tibia BM from each analyzed mouse. The absolute number of IMMs was significantly reduced only at late AML (90% reduction at t4 only and no significant loss at earlier timepoints, Fig. 2E), while that of HSMs was already drastically reduced from t2 (over 50% reduction, Fig. 2F), and the IMM:HSM ratio increased (from 20:80 in control to 45:55 at t4) as AML infiltration progressed (Fig. 2G). Together, these results revealed that the two macrophage populations present in the BM have different dynamics during AML progression. The prevalence of IMMs and rapid reduction of HSMs in AML raised the question whether IMMs may be supporting AML growth.

To understand whether IMMs and HSMs play a role in AML growth, we aimed to deplete BM macrophages using the CD169-DTR transgenic mouse model. This model was previously reported for its ability to selectively deplete F4/80+ BM macrophages after 48 hours of diphteria toxin (DT) treatment (39). In our hands, even extensive DT treatment led to a near three-fold reduction in HSMs (from 1.35×10^6^ HSM in one leg in CD169-DTR-mice to 5×10^5^ in CD169-DTR+) but did not affect IMMs (Fig. 2H-J). This may be due to differential expression levels of CD169 by IMMs and HSMs (Supplementary Fig. S2C). We injected AML cells into CD169-DTR mice after the first two daily DT injections, continued to inject DT twice a week and analyzed BM samples at a number of time points (Fig. 2K). Interestingly, AML grew significantly faster in terms of both percentage and absolute numbers of cells in the BM of DT treated CD169-DTR recipients (Fig. 2L and M). Further BM analysis revealed that in DT treated mice not only HSMs were depleted but IMMs were significantly increased, leading to a significant switch in the IMM:HSM ratio (Fig. 2N-P). The increase in IMMs was specific to DT treated, AML burdened mice, and was not induced by DT treatment alone (Fig. 2J). Hence, the presence of IMMs and absence of HSMs correlated with accelerated AML progression. These data suggest that IMMs may favor AML growth.

### IMMs present a unique efferocytic transcriptional profile

To obtain a broad and unbiased understanding of the transcriptional signatures characterizing HSMs and IMMs during BM homeostasis and AML growth, we performed single-cell RNA sequencing (scRNA-seq) on sort purified F4/80+ BM macrophages from two healthy control mice and two mice with early (15%) AML infiltration (Fig. 3A). This infiltration level was chosen at a stage because flow cytometry had identified IMMs enrichment, but no severe loss of either macrophage population had occurred yet (Fig. 3A). Over 9,000 and 10,000 cells passed quality control in control and AML samples respectively, and a median of over 2,400 genes were detected per cell (Supplementary Fig. S3A). UMAP analysis of the merged data from control and AML samples confirmed macrophages identity based on their expression of an established macrophage gene signature (*Csf1r, Cd68, Cd74* were the top expressed genes) (40), and divided them into 6 distinct macrophage clusters (Supplementary Fig. S3B and S3C). All clusters were highly conserved in both control and AML samples (Fig. 3B), with less than 50 genes being differentially expressed between control and AML samples in each cluster (Supplementary Fig. S3D and S3E). This suggested that macrophages retain their transcriptional makeup and functions at early stages of disease. Interstingly, cluster 2 expressed high levels of *Adgre1, Sirpa* and *Cd274,* encoding for proteins used for the phenotypic identification of IMMs by flow cytometry (F4/80, SIRPα and PDL-1 respectively) (Fig. 3C). In addition, clusters 0, 1 and 2 appeared enriched in the AML sample, with cluster 2 showing the strongest enrichment (Fig. 3D). Cluster 2 enrichment in the AML samples was therefore consistent with the enrichment of IMMs observed by flow cytometry. Analysis of differentially expressed genes by each cluster (Fig 3E) identified clusters 0 and 1 as monocytes and monocyte/macrophages (expressing high levels of *Nr4a1, Apoe, Ly6c2* and *Ccr2*), likely representing monocyte-derived macrophages at earlier stages of differentiation. Cluster 2 cells expressed the highest levels of genes involved in efferocytosis, such as *C1qa*, *C1qb*, *C1qc*, *Axl*, *Nr1h3* and *Mertk*. Cluster 2 cells also expressed genes involved in antigen processing and presentation, however expression of these genes was strongest in cluster 3 cells, which did not express efferocytosis markers. Cluster 4 did not show very strong DEGs. Cluster 5, while expressing high levels of the genes involved in efferocytosis, had the highest expression of genes involved in iron-recycling such as *Hmox1, Slc40a1, Spic, Hebp1* and *Trf* as well as, notably, of *Vcam1*. These data revealed that, at the molecular level, HSMs (clusters 0,1,3,4,5) are a heterogeneous population, consistent with our earlier spectral flow cytometry analyses, and include an iron-recycling macrophage population likely involved in erythropoiesis. By contrast, IMMs (cluster 2) are an efferocytic macrophage population. Efferocytosis, the process by which apoptotic cells are cleared during homeostasis and avoid inflammatory responses, can create an immune suppressive microenvironment in tumors (41). This prompted the query whether IMMs may efferocytose AML cells, contributing towards the development of an immunosuppressive microenvironment.

**Figure 3:**
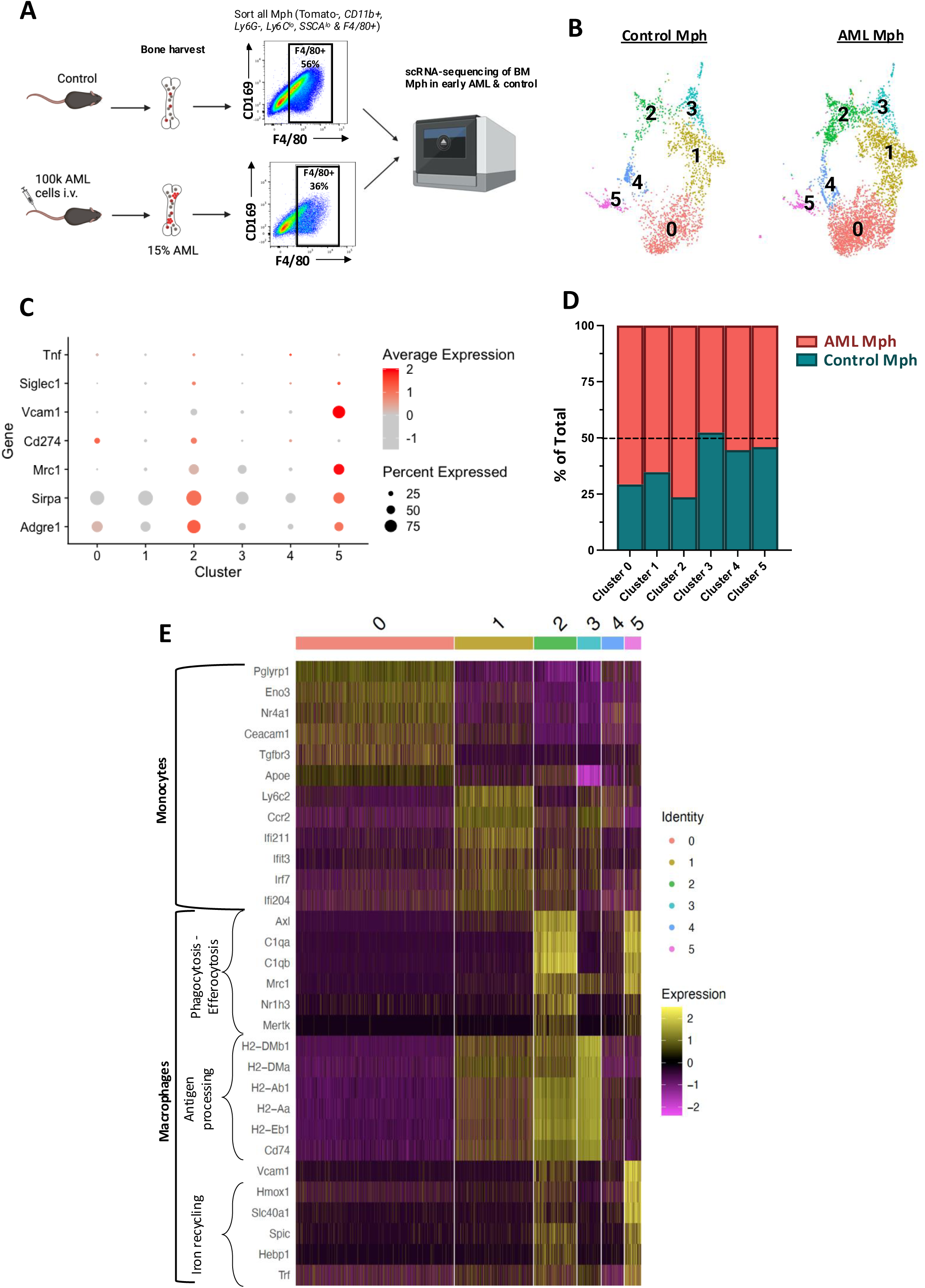
scRNA-seq identifies IMMs as an efferocytic macrophage cluster. **(A)** Bone marrow macrophages were FACS purified from naïve (*n* = *2*) or AML burdened mice (*n* = 2 per group; early AML = 17% BM infiltration) and subjected to 10X Genomics scRNA-seq. **(B)** UMAP of macrophage clusters in control and early AML bone marrow (n = 9856 individual cells for control and n = 10389 individual cells for AML). Data clustering generated 6 different clusters in both samples. **(C)** Dot Plot of the expression of genes identified by flow cytometry to identify IMMs and HSMs. **(D)** Proportion of macrophages from control and AML sample in each cluster. **(E)** Heatmap of the most differentially expressed genes for each cluster.

### IMMs efferocytose AML cells *in vitro* and *in vivo*

To uncover whether AML cells may be uptaken by macrophages, CD169-YFP mice were injected with Tomato+ AML cells and their calvaria imaged when overall BM AML infiltration was between 5-15%. High-resolution time-lapse recording revealed that CD169-YFP+ macrophages were capable of uptaking Tomato+ AML cells: this process took 10 minutes to complete, and Tomato signal could be detected inside the receiving macrophages for at least the following two hours (Fig. 4A and Supplementary Movie S4A). We quantified this process by measuring the Tomato channel mean signal intensity inside macrophages and, because autofluorescence is known to be high in macrophages, we used macrophages from control healthy mice as reference for baseline signal in the Tomato channel. We found Tomato signal in macrophages to be significantly higher in AML burdened mice (Fig. 4B). Moreover, Tomato signal in macrophages from AML burdened mice was lower than that of AML cells, confirming we were not measuring Tomato signal from adjacent AML cells (Supplementary Movie 1). To distinguish IMMs from HSMs, fluorophore conjugated anti-SIRPα antibody was injected into the mice prior to imaging, and YFP+ macrophages were segmented and classified as SIRPα high or low (Fig. 4C and Supplementary Fig. S4B) for putative IMMs (pink outlines) and HSMs (yellow outlines), respectively. Next, we analyzed the Tomato signal within each subpopulation, and we found it to be significantly higher in macrophages expressing high levels of SIRPα, or putative IMMs (Fig. 4D). These experiments provided evidence that macrophages uptake AML cells *in vivo*.

**Figure 4:**
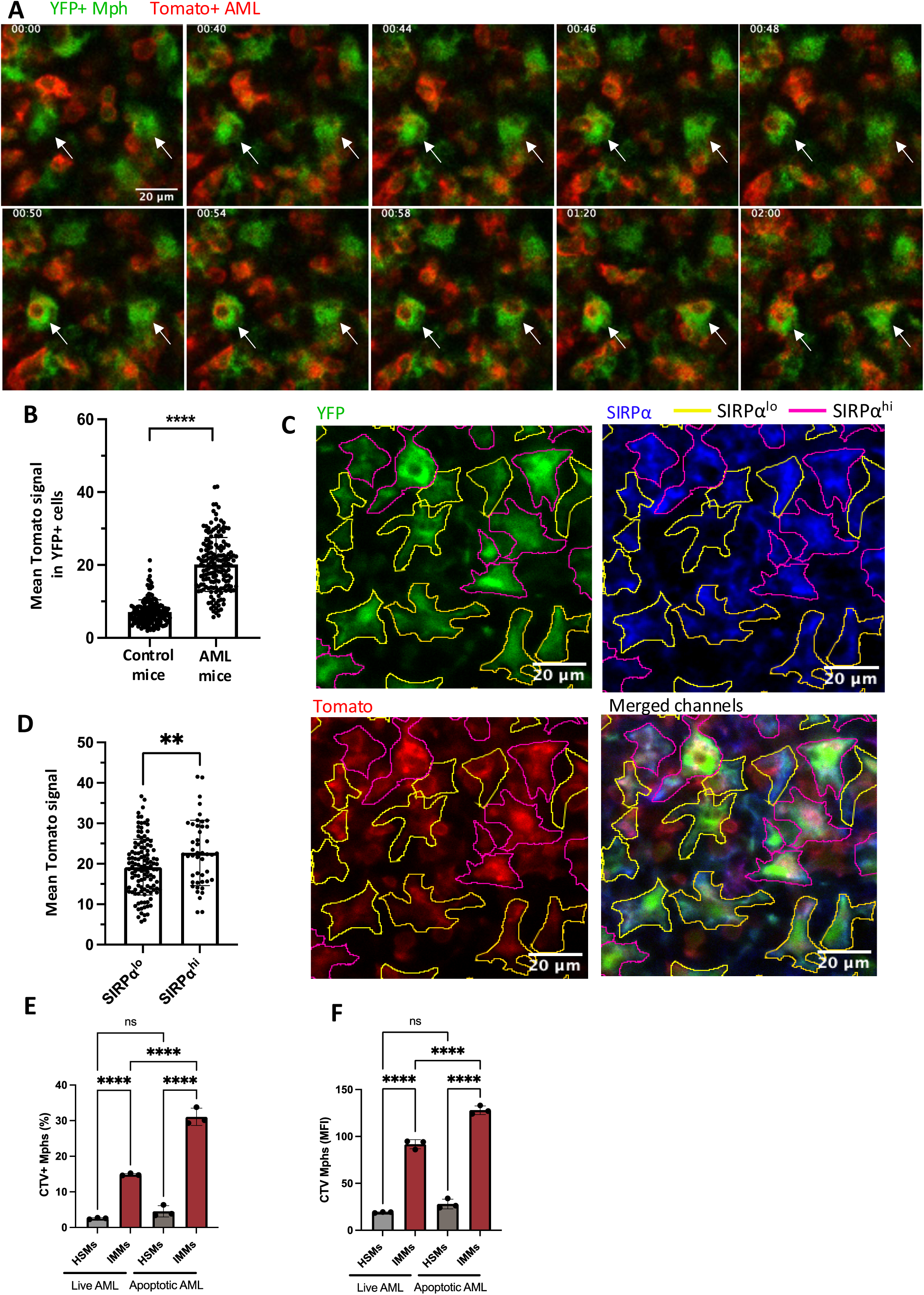
IMMs efferocytose AML cells *in vivo* and *in vitro*. **(A)** Selected time frames from representative calvarium BM two-hour timelapse of CD169-YFP mice injected with 100,000 Tomato+ AML cells and imaged at early AML stage when the BM infiltration was <15%. White arrows highlights macrophages uptaking AML cells. **(B)** Quantification of the mean Tomato signal inside CD169-YFP+ macrophages from control (n = 171 macrophages) and AML (n = 174 macrophages) mice. **(C)** Representative image of SIRP⍺ staining combined with CD169-YFP and Tomato signal. Cells were classified as SIRP⍺^lo^ (yellow outline) and SIRP⍺^hi^ (magenta outline) based on the mean intensity of the SIRP⍺ signal located inside the cell. **(D)** Quantification of the mean Tomato signal inside SIRP⍺^lo^ and SIRP⍺^hi^ CD169-YFP+ macrophages AML mice (n = 120 SIRP⍺^lo^ macrophages and n = 46 SIRP⍺^hi^ macrophages). **(E-F)** AML uptake assay of IMMs and HSMs cultured with CTV labelled, live and apoptotic AML cells. **(E)** Percentage of CTV+ macrophages. **(F)** Mean fluorescence intensity of CTV in macrophages (n = 3 donor mice, representative of 3 independent experiments). The data are presented as means ± SD. \*\**P* < 0.005; \*\*\*\**P* < 0.0001. The *P* values were determined by unpaired two-tailed Student’s *t*-test (B-D) or two-way ANOVA with post-hoc Bonferroni correction (E-F).

To further characterize the nature of AML cells’ uptake by IMMs, we set up a series of assays using IMM and HSM cell populations *in vitro*. HSMs were directly sort purified from healthy donor mice and cultured as such, with no effects on their phenotype upon 48 hours of culture (Supplementary Fig. S5A, top panel). When IMMs were sort-purified, they too maintained their phenotype in culture (Supplementary Fig. S5A, middle panel). However, IMMs are rare *in vivo* and it is challenging to obtain them in sufficient numbers for culture-based assays.

Therefore, we generated high numbers of IMMs *ex vivo* instead. We sort-purified Ly6C^high^ BM monocytes and cultured them for 48 hours, at which point flow cytometry analysis indicated the cultured cells had lost expression of Ly6C and gained high levels of SIRPα and PD-L1, hence becoming a population highly enriched for IMM-like cells (Supplementary Fig. S5A, bottom panel and Supplementary Fig. S5B and S5C). The fraction of differentiated IMMs in the culture was not affected by the addition of AML cells at the beginning of the culture (Supplementary Fig. S5D). IMMs and HSMs were co-cultured with Cell Trace Violet (CTV)-labelled live or apoptotic AML cells and the uptake of AML cells was measured through detecting CTV signal in macrophages by flow cytometry. The uptake of AML cells by IMMs was significantly higher compared to that of HSMs in both live and apoptotic AML conditions, measured as both the percentage of CTV+ macrophages (Fig. 4E) and CTV MFI (Fig. 4F). Consistent with our hypothesis that IMMs uptake AML cells via efferocytosis, in the presence of apoptotic AML cells, 30% of IMMs but only 5% of HSMs internalized AML cells (Fig. 4E). Moreover, CTV signal in IMMs was significantly higher when they were co-cultured with apoptotic AML cells, but this did not occur for HSMs (Fig. 4F).

Together, these results validated our transcriptomics analysis and confirmed IMMs as an efferocytic macrophage population, a function observed both *in vivo* and *in vitro*. Efferocytic macrophages have been associated in solid cancers with suppression of T cells’ cytotoxicity, resulting in a pro-tumoral effect (42). Consequently, these findings posed the question whether IMMs may promote AML growth through T cell inhibition.

### IMMs promote AML growth by affecting T cells

To test whether IMMs were responsible for the increase in AML growth observed in the CD169-DTR mouse model, we sought to deplete IMMs *in vivo*. Based on our *in vitro* assays that indicated IMMs to be more actively efferocytic cells than HSMs, we took advantage of clodronate liposomes, a drug that induces apoptosis following liposome internalization (43). Mice were injected with clodronate or control PBS liposomes every four days, and with AML cells the day after the first liposomes injection (Fig. 5A). PBS and clodronate liposome treated, AML burdened mice had the same amount of HSMs, while clodronate liposomes treatment prior to and during AML growth was found to significantly deplete IMMs (Fig. 5B-C). As a result, the IMM:HSM ratio was reduced in AML burdened, clodronate liposome treated mice compared to PBS treated controls (Fig. 5D). This change in the macrophage compartment was associated with a significant reduction in AML growth in both BM and peripheral blood of clodronate treated mice (Fig. 5E-F and Supplementary Fig. S6A-B).

**Figure 5:**
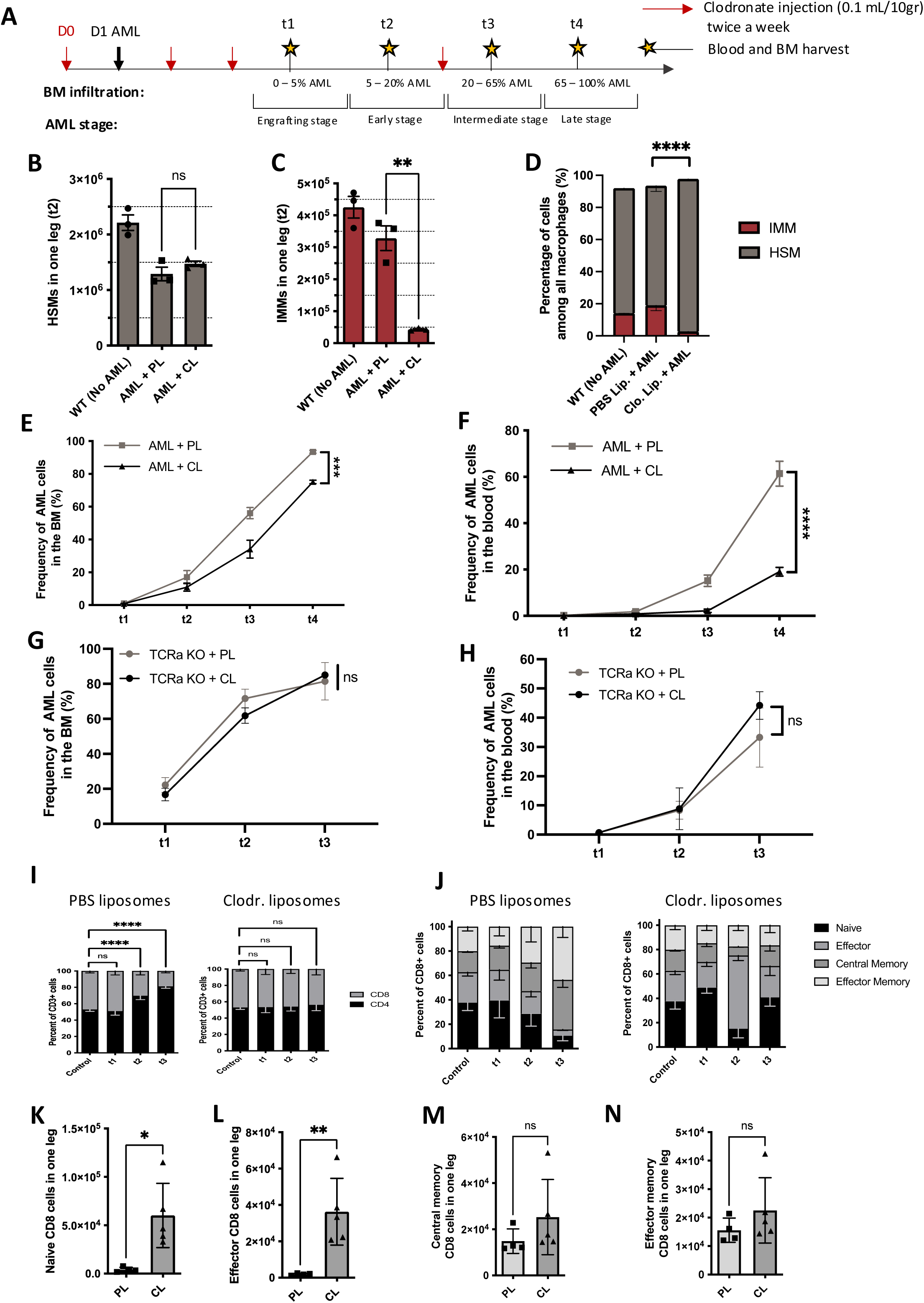
IMMs promote AML growth by inhibiting T cells. **(A)** Schematic of clodronate treatment. **(B-C)** Absolute number of BM HSMs **(B)** and IMMs **(C)** at t2 in control (no AML) and AML PBS liposome treated (PL) or clodronate liposome (CL) treated mice with AML. **(D)** Proportion of IMMs and HSMs among all BM macrophages in the mice shown in B and C. **(E-F)** Percentage of AML cells in the BM (E) and blood (F) of PBS liposome treated or clodronate liposome treated mice with AML (n = 3 mice at t1 and t2 and 4 mice at t3 and t4). **(G-H)** AML growth in BM (G) and blood (H) of TCR⍺KO mice treated with PBS liposome or clodronate liposomes. n = 5 mice per group, per timepoint. (I) Proportion of CD8+ and CD4+ cells within CD3+ cells in PBS liposome (left) or clodronate liposome (right) treated WT mice injected with AML cells. **(J)** Proportion of naïve, effcctor, central memory and effector memory CD8+ T cells in PBS liposome (left) or clodronate liposome (right) treated mice. **(K-N)** Absolute numbers of naïve (K), effector (L), central memory (M) and effector memory (N) CD8+ T cells in the BM of WT mice from I-J, at time point t3. The data are presented as means ± s.e.m. \*\**P* < 0.005; \*\*\**P* < 0.0005 \*\*\*\**P* < 0.0001. The *P* values were determined by two-way ANOVA with post-hoc Bonferroni correction.

Macrophages are known to be strong regulators of T cell activation. To evaluate whether IMMs were supporting AML growth by acting on T cells we took advantage of the TCRα knock out (KO) mouse model. These mice lack T cells but have normal levels of all other immune cells including macrophages (Supplmentary Fig. S6C) (19), therefore if the effect of IMMs on AML was T cells dependent, then in these mice clodronate liposomes should not impact AML growth. Hence, we administered clodronate liposomes to TCRα KO mice, again starting treatment on the day prior to AML cells injection. As expected, AML grew at similar speed in both the BM (Fig. 5G) and the blood (Fig. 5H) of TCRα KO treated and control mice, confirming that IMMs are supportive of AML through their interaction with T cells.

To test whether IMMs were capable of directly inhibiting T cells, we co-cultured IMMs and HSMs with pre-stimulated, CTV-labelled T cells *in vitro* and measured T cell number and cell divisions as indicators of their activation. IMMs strongly inhibited T cells’ proliferation in a concentration-dependent manner (Supplementary Fig. 6D-E). Finally, to confirm that IMMs supported AML cells growth *in vivo* by affecting T cells, we evaluated T cell subsets in AML burdened, PBS and clodronate treated mice using the well-established markers CD44 and CD62L. Interestingly, clodronate treated mice had a significant enrichment of CD8+ T cells, and especially the naïve and effector subsets T cells (Fig. 5I-J). The effect was sufficiently strong that the absolute numbers of these two T cell types were significantly increased in the BM of clodronate treated mice (Fig. 5K-N).

Taken together, all data shown so far identified IMMs as a pro-tumoral, immunosuppressive, efferocytic macrophage population, supportive of AML cells through T cell inhibition that leads to loss of effector CD8 T cells. This led us to inquire whether equivalent cells to murine IMMs may also be present in AML patients.

### IMMs are increased in fast relapsing disease

All our murine experiments indicated a pivotal role for IMMs in supporting AML growth at the early stages of BM infiltration, however the equivalent AML stages in humans are pre-clinical, and therefore not readily accessible for investigation. Instead, human samples are typically available at diagnosis, when BM infiltration is usually high, and at varying times following induction chemotherapy. We reasoned that initiation of relapse post-chemotherapy is another AML stage where AML cells are rare within the BM, and IMMs may again support AML growth. To test whether IMMs would be present following chemotherapy treatment in our murine model, we injected mice with AML cells, waited until the PB infiltration was >10%, a marker of BM infiltration >60% (19), treated them with an established cytarabine and doxorubicin (5+3) protocol (16) and finally analyzed macrophages subsets using flow cytometry (Supplementary Fig. S7A-B). The IMM:HSM ratio post-chemotherapy was significantly higher than in control, healthy untreated mice (Supplementary Fig. S7C). As the MLL-AF9 mouse model is known to develop aggressive AML regrowth following chemotherapy treatment (16), our finding raised the question whether an equivalent IMM subpopulation of human macrophages may be associated with worse prognosis in patients.

To tackle the question whether IMMs may exist and support AML growth in humans, we analyzed BM samples collected at diagnosis and post chemotherapy from AML patients with diverse cytogenetic AML subtypes, as well as some healthy BM samples (Supplementary Table S3). Given the lack of a human homolog of the uniquely specific murine macrophage marker, F4/80, we set up a high dimensional phenotyping flow cytometry analysis to identify human BM macrophages. We adopted a similar approach to that used for the murine BM flow cytometry analysis, whereby a broad myeloid panel was generated to characterize the diversity within the human BM macrophage compartment (Supplementary Table S2). To avoid any bias, all samples were pooled and cells were clustered using a combination of t-SNE and FlowSOM algorithms(44) (Fig. 6A-B). Non-myeloid cells were excluded based on their lack of expression of all myeloid markers. Neutrophils were identified as CD66b+ and inflammatory monocytes as CD14^high^ CD206-cells. AML blasts were identified as CD117^+^ CD34^+^ cells in this analysis (Fig. 6A-B), however we also knew the overall blast infiltration measured for diagnostic purposes, and percentages were corroborated with clinical diagnostic flow data. To ensure we would exclude AML-derived macrophages from our analysis, we cross-checked the frequency of AML blasts in our analysis and in the diagnostics laboratories’ reports, and excluded the few samples where frequencies did not match. In the remaining samples, two remaining clusters of healthy myeloid cells expressed CD64, CD163, CD14, CD206, SIRPα and PD-L1, albeit at different levels (Fig. 6A-B), indicating the presence of macrophage subpopulations in humans, similarly to what we observed in mice. Interestingly, the only marker necessary to separate the two macrophage clusters was the immunosupressive marker, CD206 (Fig. 6A-B), previously identified through CIBERSORT to be an important prognostic factor for AML patients(15). A simple gating strategy could then be used to identify these two macrophage populations based on CD206 expression (Supplementary Fig. S8) (45).

**Figure 6:**
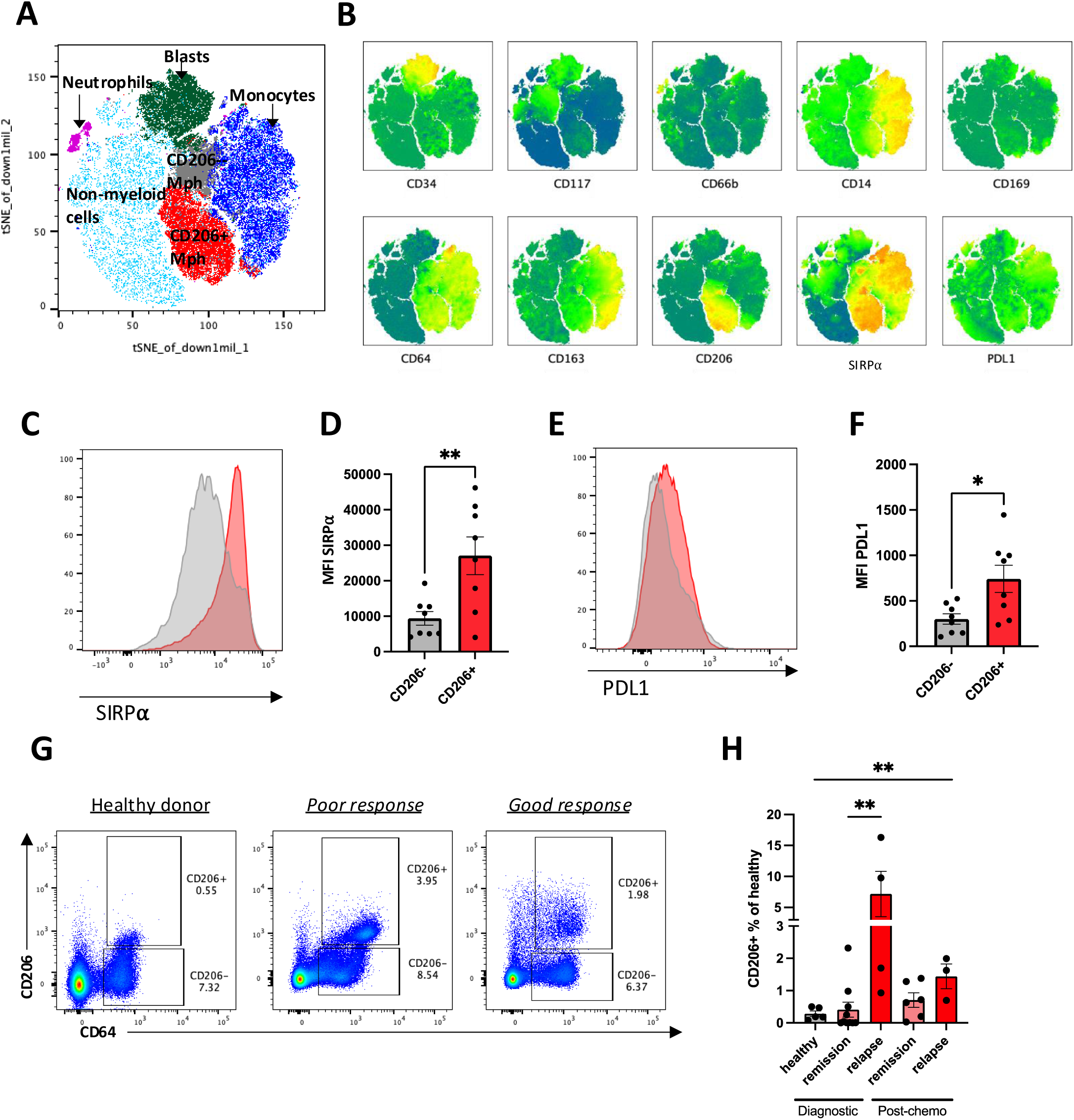
IMMs repopulate the BM after chemotherapy treatment and are increased in AML patients with recurrent disease. **(A)** Proportion of IMMs and HSMs among all macrophages in the BM of control and AML, chemotherapy treated mice (n = 3 mice per group, representative of 2 independent experiments). **(A)** t-distributed Stochastic Neighbor Embedding (t-SNE) of all single, live cells from control, diagnosis and post-chemotherapy AML BM patient samples (n= 2 controls, 6 diagnosis and paired post-chemotherapy AML). **(B)** Expression of CD34, CD117, CD66b, CD14, CD169, CD64, CD163, CD206, SIRP⍺ and PD-L1 in all live cells. **(C-D)** Histogram **(C)** and quantification **(D)** of the expression of SIRP⍺ in CD206+/- macrophages. **(E-F)** Histogram **(E)** and quantification **(F)** of the expression of PD-L1 in CD206+/- macrophages. **(G)** Representative dot plots showing relative abundance of CD206 +/- macrophages in healthy donors, good response and poor response samples. Patient samples shown here were collected post-chemotherapy. **(H)** Proportion of CD206+/- macrophages among all healthy cells in control, diagnosis and post-chemotherapy AML BM patient samples, separated by outcome. The data are presented as means ± s.e.m. \**P* < 0.05; \*\**P* < 0.005; \*\*\**P* < 0.0005 \*\*\*\**P* < 0.0001. The *P* values were determined by unpaired two-tailed Student’s *t*-test (A, E, G) or two-way ANOVA with post-hoc Bonferroni correction (I).

To correlate these clusters with the murine BM macrophage populations, we evaluated the co-expression of SIRPα, PD-L1 and CD206+. Human CD206+ macrophages were found to express significantly higher levels of SIRPα (Fig. 6C and D) and PD-L1 (Fig. 6E and F), consistent with the murine IMMs’ phenotype. Conversely, we assessed expression of CD206 in murine IMMs and HSMs in healthy mice. Strikingly, HSMs did not express CD206, while a clear fraction of IMMs were positive for this marker (Supplementary Fig. S8B-C). In addition, the fraction of IMMs expressing CD206 increased significantly with AML growth, reaching 50% at late AML stage (Supplementary Fig. S8D).

To test whether IMMs/CD206+ human macrophages may be associated with worse prognosis in patients, we measured the frequency of CD206+ macrophages among healthy cells in BM samples from healthy donors and from AML patients that maintained remission for over one year from end of chemotherapy or after allogeneic hematopoietic stem cell transplantation (classified as good response) or not (classified as poor response). Consistent with other studies, IMMs frequency within healthy cells was highly variable in diagnostic samples, however was significantly higher in samples from patients with poor outcome compared to those with good outcome (Fig. 6G and H). In addition, in samples collected after chemotherapy, the frequency of CD206+ macrophages in patients that went into and maintained remission was similar to that of healthy donors, but was significantly higher in patients that relapsed early (Fig. 6G and H).

## Discussion

Acute myeloid leukemia (AML) maintains a bleak prognosis due to its propensity for relapse post-initial chemotherapy response(46–49). Furthermore, despite the transformative potential of immunotherapies in cancer treatment, AML exhibits markedly limited response rates. The mechanisms driving immune evasion and consequent suboptimal responsiveness to immunotherapies in AML are not yet fully understood, because it is experimentally challenging to model physiological interactions between AML, innate and adaptive immune cells. The syngeneic, conditioning-free *in vivo* murine model of AML we used affords unique insights into the different macrophage populations within the BM, their interactions with AML cells and T cells, and the resulting effect on AML growth.

Using the CD169-YFP macrophage reporter mouse line, spatial analysis of the BM at early AML infiltration revealed an enrichment of YFP+ macrophages in AML foci. Given that all healthy hematopoietic cells are rapidly lost with AML(16,27), this enrichment was suggestive of a role of macrophages in promoting disease growth. High dimensional spectral flow cytometry analysis revealed a high level of heterogeneity within the macrophage population, correlating with previous identification of numerous macrophage populations associated with distinct functions within the BM(36–38,50). However, one specific sub-population of macrophages was enriched in the BM at early AML and could be distinguished based on its expression of SIRPα and PD-L1. SIRPα and PD-L1 are two known immune checkpoints(51–53), suggesting that these cells may play a role in the host’s immune response and as such this population of macrophages was named immune-modulatory macrophages (IMMs), while all other macrophages were grouped under the name of hematopoiesis-supportive macrophages (HSMs), based on their role in the regulation of the hematopoietic system.

Using the CD169-DTR mouse model, we depleted HSMs and enriched the environment for IMMs and found that AML grew faster, correlating with a pro-tumoral role for IMMs. This is consistent with a previous study that described SIRPα+ macrophages as responsible for T cell suppression, high phagocytosis of tumor cells and reduced survival in patients(54). scRNA-seq analysis of all BM macrophages in our AML model revealed an efferocytic function for IMMs and confirmed the iron-recycling function for a sub-population of HSMs, correlating with EBIs. Importantly, our study highlights that IMMs are present in low number in healthy BM and their molecular signature is conserved as AML grows. Intravital microscopy and *in vitro* co-culture validated that IMMs were efferocytic as they were capable of uptaking apoptotic AML cells. By preferentially depleting IMMs with clodronate liposomes, we were able to confirm their pro-tumoral function because AML grew slower in treated mice. Given the importance of efferocytic macrophages in T cell inhibition in solid tumors(42), we depleted IMMs in mice devoid of functional T cells (TCRαKO) and found that the effect IMMs had on AML was mediated by T cells. Co-cultures of T cells with IMMs validated that IMMs inhibited T cells and could therefore be confirmed as an immunosuppressive cell type. Importantly, IMM depletion led to an enrichment in naïve and effector CD8+ T cells. Further studies will be required to understand the specific and dynamic mechanisms through which AML triggers imbalances in macrophage and T cell populations. The expression of antigen-presentation machinery genes by efferocytic IMMs, albeit at lower levels than other macrophage subpopulations, raises the question whether this antigen presentation may be a mechanism through which IMMs engage T cells. Interestingly, a recent study showed that inhibition of Axl, a known efferocytic receptor expressed by macrophages, stimulated host-versus-leukemia immunity and eradicated leukemia in a murine B cell acute lymphoblastic leukaemia model (55). Our results are in line with these findings as they highlight the major role efferocytic macrophages play in AML development and the need for their targeting to improve the efficacy of immunotherapies in AML.

Whether HSMs and/or IMMs are tissue resident macrophages or monocyte-derived remains an open question. Whichever the answer to this question may be, the important finding of our study is that targeting macrophages affects AML growth. This is relevant for recent initial trials using CD47 blocking antibodies(56), and our work provides a further mechanism whereby blocking efferocytosis and ideally enhancing both antigen presentation and immune activation would be the most successful intervention.

Similarly to mice, we could also identify two populations of macrophages in humans, distinguishable based on their CD206 expression. The macrophage population in mice and human may not be totally overlapping, nevertheless CD206 expression was found only in IMMs in mice, suggesting that the human CD206+ macrophages are equivalent to murine IMMs. At diagnosis, poor outcome patients had higher frequencies of CD206+ cells than good outcome patients. Post-chemotherapy, the frequency of CD206+ cells was increased in patients with poorer outcomes, while it was similar in healthy donors and in patients who responded to chemotherapy and remain in ongoing remission. Consistent with our finding, in multiple myeloma, patients with a high ratio of M2/M1 macrophages were shown to have lower response rates to therapy and reduced progression free survival and overall survival(12). In addition, through sequencing, several studies have shown that CD206 is associated with a poor survival in humans with AML(9,15). However, these studies did not elucidate the mechanism behind this result. Interstingly, a recent study reported the abundance of CD206+ CD163+ pro-inflammatory macrophages to correlate with poor outcome (57). Our findings highlight the heterogeneity of macrophage populations and their complex interactions with multiple cell types, with different subpopulations likely acting at different times or within different BM region to drive disease progression. Based on these results, we propose an immune evasion mechanism whereby an immunossupressive, efferocytic, pro-tumoral macrophage population, IMMs, expressing CD206, uptake apoptotic AML cells and inhibit T cells resulting in AML growth. These findings justify the poor survival associated with CD206 expression and highlight the need to target macrophages as well as T cells for immunotherapies to be efficient.

T cells responses have been identified in murine AML models and patients, however are known to eventually fail (58, 59). By exploring the heterogeneity of BM macrophages in mice and humans we are able to propose a mechanism underpinning such failure. Our findings indicate that targeting SIRPα+ PDL-1+ macrophages, either by reducing their numbers or by remodelling their function may be a valuable therapeutic avenue to improve AML outcome.

## Acknowledgments

We thank Muzliffah Haniffa, Ilaria Malanchi and Marina Botto for enriching discussions on macrophage biology. We thank the Sir Francis Crick (Crick) and Imperial College London genomics facilities for the single cell transcriptomics planning and work, animal facilities and flow cytometry facilities for technical support.

FB and CM were funded by Fondation Alcea; CLC by CRUK (Programme Foundation Award C36195/A26770, Fondation Alcea, and the Wellcome Trust (Investigator Award 212304/Z/18/Z). MLRH was funded by Wellcome Trust PhD studentship 105398/Z/14/Z and Sir Henry Wellcome Postdoctoral Fellowship 224055/Z/21/Z; SGA by a CRUK PhD studentship C36195/A27830; RJB by CRUK Clinician Scientist Fellowship RCCFEL/100017. CA received funding from the ISREC Foundation.

## Supplementary Materials

**Supplementary Figure 1: Morphology of macrophages in AML early stage mice**

CD169-Cre x R26-YFP mice were injected with 100,000 Tomato+ AML cells and imaged at the early stage of AML (t2). YFP+ macrophages were manually segmented and analyzed. **(A-C)** Circularity **(A),** perimeter **(B)** and area **(C)** of YFP+ macrophages per ROI classified according to the fraction of AML in each ROIs. n= 1378 macrophages in a ‘single cells’ ROIs, 702 macrophages in b ‘small clusters’ ROIs, 730 macrophages in c ‘large clusters’ ROIs from 3 mice. The data are presented as violin plots, median is represented with black dotted lines and 1^st^ and 3^rd^ interquartiles are shown in grey dotted lines. \*\**P* < 0.005; \*\*\**P* < 0.0005 \*\*\*\**P* < 0.0001. The *P* values were determined by unpaired two-tailed Student’s *t*-test.

**Supplementary Figure 2: Gating strategy and characterization of macrophage subpopulations in murine BM**

**(A)** Manual gating strategy used to identify macrophages in the BM. **(B)** UMAPs displaying the expression of CD11b, Ly6G, Ly6C, SSC-A, F4/80, Tim4, iNOS, MerTK, VEGF-A, CD86, LYVE1, RELM⍺, Arg1, TGFß and CD115 in macrophages pooled from control and early AML BM (n= 3 controls and 3 early AML mice). **(C)** Representative histogram of CD169 expression in IMMs and HSMs in the BM.

**Supplementary Figure 3: scRNA seq macrophage analysis of macrophages**

**(A)** Number of cells that passed quality control and median number of genes detected per cell per sample analysed by scRNA seq. **(B)** Merged UMAP of all macrophage clusters from control and early AML bone marrow combined samples. **(C)** Dot Plot of the expression of the core macrophages genes signature identified by E. Gautiar et al.(40) **(D)** Overlap of control and early AML samples data. **(E)** Number of DEGs between control and early AML clusters.

**Supplementary Figure 4: Macrophage efferocytose AML cells in vivo**

**(A)** Quantification of the mean Tomato signal inside CD169-YFP+ macrophages (n = 174) and AML cells (n = 132). Bars reprsent means ± s.e.m. Each dot is one cell analyzed. \*\*\*\**P* < 0.0001. The *P* value was determined by unpaired two-tailed Student’s *t*-test. **(B)** Histogram of the expression of SIRP⍺ in YFP+ macrophages. The threshold between SIRP⍺^lo^ and SIRP⍺^hi^ was determined based on the proportion of known SIRP⍺^hi^ cells identified by FACS and the distribution of the histogram.

**Supplementary Figure 5: Ly6C^hi^ monocytes differentiate into IMMs**

**(A)** Manual gating strategy used to identify HSM/IMMs following 48 hours in culture of sorted HSMs, IMMs and Ly6C^hi^ cells. **(B)** Ly6C mean fluorescence intensity of Ly6C^hi^ sorted cells after 24 and 48 hours in culture. **(C)** Proportion of IMMs following 48 hours in culture of sorted Ly6C^hi^ cells, HSMs, and IMMs. **(D)** Proportion of IMMs following 48 hours in culture of Ly6C^hi^ cells, cultured alone or with AML cells. Data are presented as means ± s.e.m. \*\*\*\**P* < 0.0001. ns: not significant. The *P* values were determined by unpaired two-tailed Student’s *t*-test (B, D) or two-way ANOVA with post-hoc Bonferroni correction (C).

**Supplementary Figure 6: IMMs support AML growth by inhibiting T cells**

**(A-B)** Absolute number of AML cells in the BM **(A)** and blood **(B)** of PBS liposome treated or clodronate liposome treated mice with AML (n = 3 mice at t1 and t2 and 4 mice at t3 and t4). **(C)** Absolute numbers of HSMs (left) and IMMs (right) per combined femur and tibia (leg) of wild type and TCRα KO mice. n= 5 mice/group. The *P* values were determined by unpaired two-tailed Student’s *t*-test and were >0.05. **(D-E)** Co-cultures of HSMs or IMMs and stimulated, CTV labelled T cells. **(D)** Representative example of T-cell proliferation as measured by dilution of CTV. Purified T cells were labelled with CTV, activated *in vitro* by CD3/CD28 and cocultured with the indicated ratio of IMMs (left) and HSMs (right) for 60 h. Live, single CD3+ cells are shown. **(E)** Division index (left) and total T cell count (right) of T cells cultured alone or in the presence of IMMs or HSMs at a ratio of 1:0.4. Data are presented as means ± s.e.m. \**P* < 0.05; \*\**P* < 0.005; \*\*\*\**P* < 0.0001. The *P* values were determined by unpaired two-tailed Student’s *t*-test (A-C) or two-way ANOVA with post-hoc Bonferroni correction (E-F).

**Supplementary Figure 7: IMMs are enriched post-chemotherapy**

**(A)** Representative experimental plan. Mice were injected with 100,000 MLL-AF9 AML cells i.v., and their PB was monitored. When AML infiltration was >10% (day 20 in the specific example), chemotherapy was administered for 5 days. PB continued to be monitored and 6 days post-chemotherapy the experiment was terminated and BM was harvested and analyzed. **(B)** Longitudinal flow cytometry analysis indicating AML PB infiltration decreasing after the first dose of chemotherapy and starting to increase in some animals by the time BM was analyzed. **(C)** Proportion of IMM and HSM subpopulations of BM microphages in healthy, untreated mice and in mice that received AML cells and chemotherapy as shown in (A). Data are presented as means ± s.e.m. \*\*\*\**P* < 0.0001. The *P* values was determined by unpaired two-tailed Student’s *t*-test.

**Supplementary Figure 8: Human CD206+ macrophages are similar to murine IMMs.**

**(A)** Manual gating strategy used to identify blasts, neutrophils, monocytes and CD206+/- macrophages in AML patients BM samples. **(B)** Representative histogram of the expression of CD206 in IMMs and HSMs in healthy control mice. **(C)** Proportion of CD206+ HSMs and IMMs (n = 8). **(D)** Proportion of IMMs expressing CD206 throughout AML progression (n = 3 - 8 mice per timepoint). Data are presented as means ± s.e.m. \**P* < 0.05; \*\**P* < 0.005; \*\*\**P* < 0.0005 \*\*\*\**P* < 0.0001. The *P* values were determined by unpaired two-tailed Student’s *t*-test (C) or two-way ANOVA with post-hoc Bonferroni correction (D).

**Supplementary Table S1: Murine myeloid antibodies used in this study.**

Conjugated fluorophore, clone, dilution and source of each antibody used in this study to analyze murine samples.

**Supplementary Table S2: Human antibodies used in this study.**

Conjugated fluorophore, clone, dilution and source of each antibody used in this study to analyze human samples.

**Supplementary Table S3: Details of AML patient samples used.**

Bone marrow samples were collected at diagnosis or following blood count recovery following the first cycle of induction chemotherapy (all within 4-6 weeks after start of chemotherapy). Grey background: good response. White background: poor response. Treatments: DA = Daunorubicin + Cytarabine; HiDAC = High dose Cytarabine; Go = Gemtuzumab Ozogamicin; FLA-Ida = Fludarabine + Cytarabine + Idarubicin; MA = Mitoxantrone + Cytarabine; FLA = Fludarabine + Cytarabine; Bu/Cy ASCT = Busulfan + Cyclophosphamide Allogeneic Stem Cell Transplant; EM = XXX; FLAG = Fludarabine + Cytarabine + G-CSF + Idarubicin; IA = Idarubicin + Cytarabine; AZAVEN = Azacytidine + Venetoclax; allo = allogeneic bone marrow transplant; A = cytarabine; AZA = Azacytidine. Patients 1 to 6 were treated at UCLH; patients 7 to 15 at CHUV.

---

**Supplementary Video 1:**
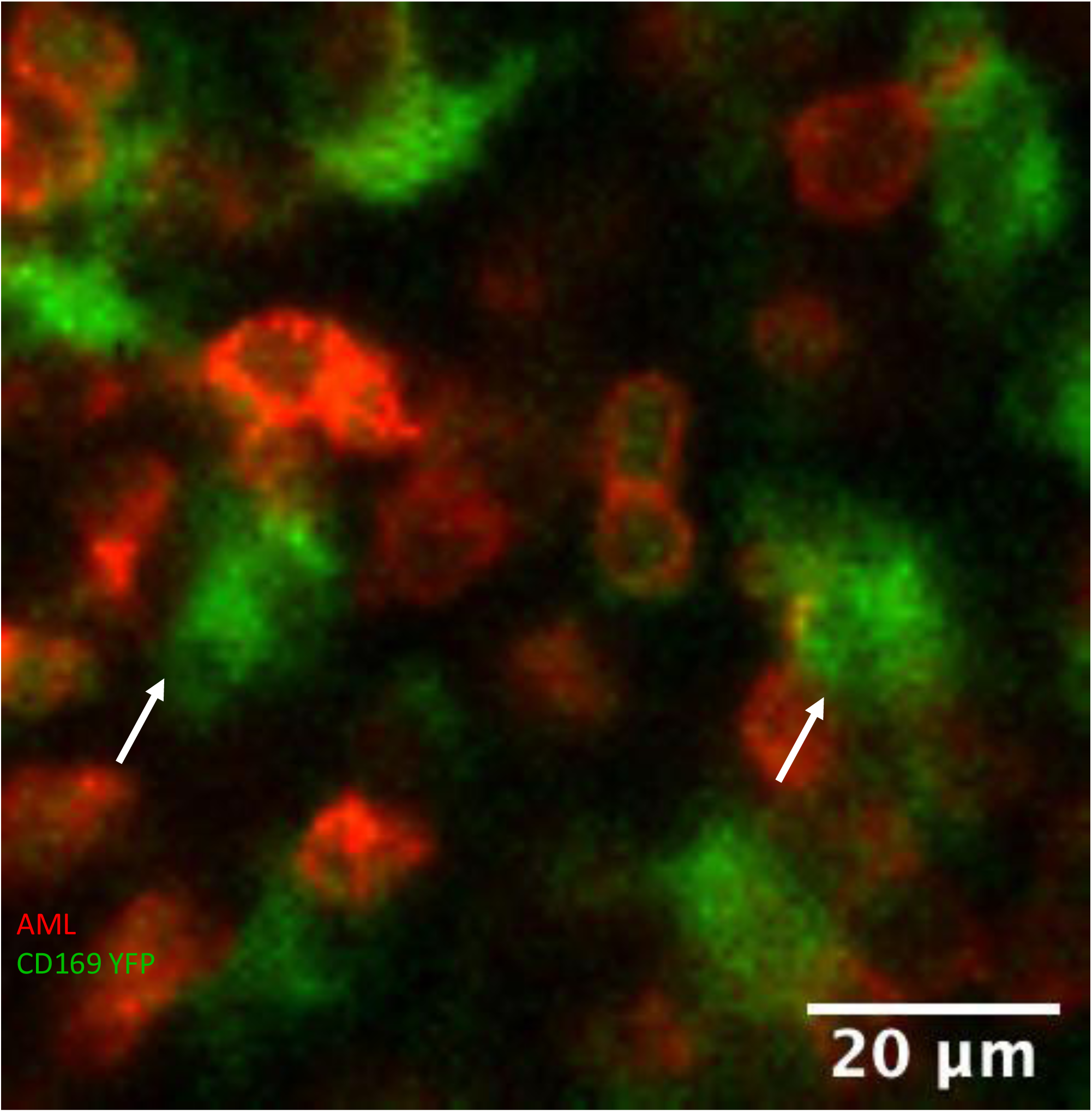
**Uptake of mTmG+ AML cells by CD169-YFP+ cells.** Representative maximum projection of two-hour long time-lapse with images acquired every two minutes from the BM of an AML burdened CD169-YFP^het^ mouse. Green: CD169-YFP+ cells, Red: AML cells (uptake of AML cells could be observed 1-2 times/ frame, 4 frames per mice were acquired, n= 3 mice).

## References

1. Bassan R, Bourquin JP, DeAngelo DJ, Chiaretti S. New Approaches to the Management of Adult Acute Lymphoblastic Leukemia. 10.1200/JCO2017773648. American Society of Clinical Oncology; 2018;36:3504–19.

2. Kantarjian H, Kadia T, DiNardo C, Daver N, Borthakur G, Jabbour E, et al. Acute myeloid leukemia: current progress and future directions. Blood Cancer Journal 2021 11:2 [Internet]. Nature Publishing Group; 2021 [cited 2022 Sep 19];11:1–25. Available from: https://www.nature.com/articles/s41408-021-00425-3

3. Sauerer T, Velázquez GF, Schmid C. Relapse of acute myeloid leukemia after allogeneic stem cell transplantation: immune escape mechanisms and current implications for therapy. Molecular Cancer 2023 22:1. BioMed Central; 2023;22:1–22.

4. Witkowski MT, Lasry A, Carroll WL, Aifantis I. Immune-Based Therapies in Acute Leukemia. Trends Cancer. Elsevier; 2019;5:604–18.

5. Jan M, Leventhal MJ, Morgan EA, Wengrod JC, Nag A, Drinan SD, et al. Recurrent genetic HLA loss in AML relapsed after matched unrelated allogeneic hematopoietic cell transplantation. Blood Adv. American Society of Hematology; 2019;3:2199–204.

6. Vago L, Perna SK, Zanussi M, Mazzi B, Barlassina C, Stanghellini MTL, et al. Loss of Mismatched HLA in Leukemia after Stem-Cell Transplantation. New England Journal of Medicine. Massachusetts Medical Society; 2009;361:478–88.

7. Toffalori C, Zito L, Gambacorta V, Riba M, Oliveira G, Bucci G, et al. Immune signature drives leukemia escape and relapse after hematopoietic cell transplantation. Nat Med. Nat Med; 2019;25:603–11.

8. Jaiswal S, Jamieson CHM, Pang WW, Park CY, Chao MP, Majeti R, et al. CD47 Is Upregulated on Circulating Hematopoietic Stem Cells and Leukemia Cells to Avoid Phagocytosis. Cell. Cell Press; 2009;138:271–85.

9. van Galen P, Hovestadt V, Wadsworth MH, Hughes TK, Griffin GK, Battaglia S, et al. Single-Cell RNA-Seq Reveals AML Hierarchies Relevant to Disease Progression and Immunity. Cell. Cell Press; 2019;176:1265-1281.e24.

10. Mass E, Nimmerjahn F, Kierdorf K, Schlitzer A. Tissue-specific macrophages: how they develop and choreograph tissue biology. Nature Reviews Immunology |. 2023;23:563.

11. Al-Matary YS, Botezatu L, Opalka B, Hönes JM, Lams RF, Thivakaran A, et al. Acute myeloid leukemia cells polarize macrophages towards a leukemia supporting state in a Growth factor independence 1 dependent manner. Haematologica. Ferrata Storti Foundation; 2016;101:1216.

12. Yang X, Feng W, Wang R, Yang F, Wang L, Chen S, et al. Repolarizing heterogeneous leukemia-associated macrophages with more M1 characteristics eliminates their pro-leukemic effects. Oncoimmunology. Taylor and Francis Inc.; 2018;7.

13. Keech T, McGirr C, Winkler IG, Levesque J-P. Macrophage Involvement in the Response of Acute Myeloid Leukaemia to Chemotherapy. Blood. American Society of Hematology; 2017;130:5069–5069.

14. Gilead Sciences. Gilead statement on discontinuation of phase 3 ENHANCE-3 study in AML [Internet]. 2024 [cited 2024 Mar 27]. Available from: http://tinyurl.com/347tjc2h

15. Xu ZJ, Gu Y, Wang CZ, Jin Y, Wen XM, Ma JC, et al. The M2 macrophage marker CD206: a novel prognostic indicator for acute myeloid leukemia. Oncoimmunology. Taylor and Francis Inc.; 2020;9.

16. Duarte D, Hawkins ED, Akinduro O, Ang H, De Filippo K, Kong IY, et al. Inhibition of Endosteal Vascular Niche Remodeling Rescues Hematopoietic Stem Cell Loss in AML. Cell Stem Cell. Cell Press; 2018;22:64-77.e6.

17. Krivtsov A V., Figueroa ME, Sinha AU, Stubbs MC, Feng Z, Valk PJM, et al. Cell of origin determines clinically relevant subtypes of MLL-rearranged AML. Leukemia. NIH Public Access; 2013;27:852.

18. Akinduro O, Weber TS, Ang H, Haltalli MLR, Ruivo N, Duarte D, et al. Proliferation dynamics of acute myeloid leukaemia and haematopoietic progenitors competing for bone marrow space. Nat Commun. Nature Publishing Group; 2018;9:1–12.

19. Mombaerts P, Clarke AR, Rudnicki MA, Iacomini J, Itohara S, Lafaille JJ, et al. Mutations in T-cell antigen receptor genes a and b block thymocyte development at different stages. Nature.; 1992; 360:225–231.

20. Hawkins ED, Duarte D, Akinduro O, Khorshed RA, Passaro D, Nowicka M, et al. T-cell acute leukaemia exhibits dynamic interactions with bone marrow microenvironments. Nature. Nature; 2016;538:518–22.

21. Zheng GXY, Terry JM, Belgrader P, Ryvkin P, Bent ZW, Wilson R, et al. Massively parallel digital transcriptional profiling of single cells. Nature Communications 2017 8:1 [Internet]. Nature Publishing Group; 2017 [cited 2022 Apr 27];8:1–12. Available from: https://www.nature.com/articles/ncomms14049

22. Dobin A, Davis CA, Schlesinger F, Drenkow J, Zaleski C, Jha S, et al. Sequence analysis STAR: ultrafast universal RNA-seq aligner. 2013;29:15–21.

23. R: A Language and Environment for Statistical Computing Reference Index The R Development Core Team. [cited 2024 Mar 9]; Available from: http://www.gnu.org/copyleft/gpl.html.

24. Butler A, Hoffman P, Smibert P, Papalexi E, Satija R. Integrating single-cell transcriptomic data across different conditions, technologies, and species Analysis. nature biotechnology VOLUME. 2018;36.

25. Subramanian A, Tamayo P, Mootha VK, Mukherjee S, Ebert BL, Gillette MA, et al. Gene set enrichment analysis: A knowledge-based approach for interpreting genome-wide expression profiles. Proc Natl Acad Sci U S A. National Academy of Sciences; 2005;102:15545–50.

26. Yu G, Wang LG, Han Y, He QY. clusterProfiler: an R package for comparing biological themes among gene clusters. OMICS. OMICS; 2012;16:284–7.

27. Pirillo C, Birch F, Tissot FS, Anton SG, Haltalli M, Tini V, et al. Metalloproteinase inhibition reduces AML growth, prevents stem cell loss, and improves chemotherapy effectiveness. Blood Adv. 2022;6.

28. Wei Q, Boulais PE, Zhang D, Pinho S, Tanaka M, Frenette PS. Maea expressed by macrophages, but not erythroblasts, maintains postnatal murine bone marrow erythroblastic islands. Blood. American Society of Hematology; 2019;133:1222–32.

29. Karasawa K, Asano K, Moriyama S, Ushiki M, Monya M, Iida M, et al. Vascular-resident CD169-positive monocytes and macrophages control neutrophil accumulation in the kidney with ischemia-reperfusion injury. Journal of the American Society of Nephrology. American Society of Nephrology; 2015;26:896–906.

30. Asano K, Takahashi N, Ushiki M, Monya M, Aihara F, Kuboki E, et al. Intestinal CD169+ macrophages initiate mucosal inflammation by secreting CCL8 that recruits inflammatory monocytes. Nature Communications 2015 6:1. Nature Publishing Group; 2015;6:1–14.

31. Camara A, Lavanant AC, Abe J, Desforges HL, Alexandre YO, Girardi E, et al. CD169+ macrophages in lymph node and spleen critically depend on dual RANK and LTbetaR signaling. Proc Natl Acad Sci U S A. National Academy of Sciences; 2022;119:e2108540119.

32. McInnes L, Healy J, Melville J. UMAP: Uniform Manifold Approximation and Projection for Dimension Reduction. 2018;

33. Duan Z, Luo Y. Targeting macrophages in cancer immunotherapy. Signal Transduction and Targeted Therapy 2021 6:1 [Internet]. Nature Publishing Group; 2021 [cited 2024 Mar 27];6:1–21. Available from: https://www.nature.com/articles/s41392-021-00506-6

34. Bisht K, McGirr C, Lee SY, Tseng HW, Fleming W, Alexander KA, et al. Oncostatin M regulates hematopoietic stem cell (HSC) niches in the bone marrow to restrict HSC mobilization. Leukemia 2021 36:2. Nature Publishing Group; 2021;36:333–47.

35. Kaur S, Raggatt LJ, Millard SM, Wu AC, Batoon L, Jacobsen RN, et al. Self-repopulating recipient bone marrow resident macrophages promote long-term hematopoietic stem cell engraftment. Blood. American Society of Hematology; 2018;132:735–49.

36. Dutta P, Hoyer FF, Grigoryeva LS, Sager HB, Leuschner F, Courties G, et al. Macrophages retain hematopoietic stem cells in the spleen via VCAM-1. Journal of Experimental Medicine. Rockefeller University Press; 2015;212:497–512.

37. Li W, Wang Y, Zhao H, Zhang H, Xu Y, Wang S, et al. Identification and transcriptome analysis of erythroblastic island macrophages. Blood. American Society of Hematology; 2019;134:480–91.

38. Li D, Xue W, Li M, Dong M, Wang J, Wang X, et al. VCAM-1+ macrophages guide the homing of HSPCs to a vascular niche. Nature 2018 564:7734. Nature Publishing Group; 2018;564:119–24.

39. Chow A, Lucas D, Hidalgo A, Méndez-Ferrer S, Hashimoto D, Scheiermann C, et al. Bone marrow CD169+ macrophages promote the retention of hematopoietic stem and progenitor cells in the mesenchymal stem cell niche. Journal of Experimental Medicine. Rockefeller University Press; 2011;208:761–71.

40. Gautiar EL, Shay T, Miller J, Greter M, Jakubzick C, Ivanov S, et al. Gene-expression profiles and transcriptional regulatory pathways that underlie the identity and diversity of mouse tissue macrophages. Nat Immunol. 2012;13.

41. Werfel TA, Cook RS. Efferocytosis in the tumor microenvironment. Semin Immunopathol [Internet]. Springer; 2018 [cited 2024 Mar 27];40:545. Available from: pmc/articles/PMC6223858/

42. Qiu H, Shao Z, Wen X, Liu Z, Chen Z, Qu D, et al. Efferocytosis: An accomplice of cancer immune escape. Biomedicine & Pharmacotherapy. Elsevier Masson; 2023;167:115540.

43. Rooijen N Van, Sanders A. Liposome mediated depletion of macrophages: mechanism of action, preparation of liposomes and applications. J Immunol Methods. Elsevier; 1994;174:83–93.

44. Van Gassen S, Callebaut B, Van Helden MJ, Lambrecht BN, Demeester P, Dhaene T, et al. FlowSOM: Using self-organizing maps for visualization and interpretation of cytometry data. Cytometry Part A. John Wiley & Sons, Ltd; 2015;87:636–45.

45. Yu YRA, Hotten DF, Malakhau Y, Volker E, Ghio AJ, Noble PW, et al. Flow cytometric analysis of myeloid cells in human blood, bronchoalveolar lavage, and lung tissues. Am J Respir Cell Mol Biol [Internet]. American Thoracic Society; 2016 [cited 2024 Mar 27];54:13–24. Available from: https://experts.arizona.edu/en/publications/flow-cytometric-analysis-of-myeloid-cells-in-human-blood-bronchoa

46. Burnett AK, Russell NH, Hills RK, Kell J, Cavenagh J, Kjeldsen L, et al. A randomized comparison of daunorubicin 90 mg/m2 vs 60 mg/m2 in AML induction: results from the UK NCRI AML17 trial in 1206 patients. Blood. Blood; 2015;125:3878–85.

47. Freeman SD, Thomas A, Thomas I, Hills RK, Vyas P, Gilkes A, et al. Fractionated vs single-dose gemtuzumab ozogamicin with determinants of benefit in older patients with AML: the UK NCRI AML18 trial. Blood. Elsevier B.V.; 2023;142:1697–707.

48. Yilmaz M, Wang F, Loghavi S, Bueso-Ramos C, Gumbs C, Little L, et al. Late relapse in acute myeloid leukemia (AML): clonal evolution or therapy-related leukemia? Blood Cancer Journal 2019 9:2. Nature Publishing Group; 2019;9:1–6.

49. Shlush LI, Mitchell A, Heisler L, Abelson S, Ng SWK, Trotman-Grant A, et al. Tracing the origins of relapse in acute myeloid leukaemia to stem cells. Nature 2017 547:7661. Nature Publishing Group; 2017;547:104–8.

50. Haldar M, Kohyama M, So AYL, Kc W, Wu X, Briseño CG, et al. Heme-Mediated SPI-C Induction Promotes Monocyte Differentiation into Iron-Recycling Macrophages. Cell. Cell Press; 2014;156:1223–34.

51. Sun C, Mezzadra R, Schumacher TN. Immunity Review Regulation and Function of the PD-L1 Checkpoint. Immunity. 2018;48:434–52.

52. Veillette A, Chen J. SIRPα–CD47 Immune Checkpoint Blockade in Anticancer Therapy. Trends Immunol. Elsevier Ltd; 2018;39:173–84.

53. Logtenberg ME, Scheeren FA, Schumacher TN. The CD47-SIRP&#x03B1; Immune Checkpoint. Immunity. 2020;52:742–52.

54. Chen gYP, Kim HJ, Wu H, Price-Troska T, Villasboas JC, Jalali S, et al. SIRPα expression delineates subsets of intratumoral monocyte/macrophages with different functional and prognostic impact in follicular lymphoma. Blood Cancer J. Nature Publishing Group; 2019;9:1–14.

55. Tirado-Gonzalez I, Descot A, Soetopo D, Nevmerzhitskaya A, Schäffer A, Kur IM, et al. Axl inhibition in macrophages stimulates host-versus-leukemia immunity and eradicates naïve and treatment-resistant leukemia. Cancer Discov. American Association for Cancer Research Inc.; 2021;11:2924–43.

56. Bouwstra R, Meerten T van, Bremer E. CD47-SIRPα blocking-based immunotherapy: Current and prospective therapeutic strategies. Clin Transl Med. Wiley-Blackwell; 2022;12.

57. Weinhauser I, Pereira-Martins DA, Almeida LY, Hilberink JR, Silveira DRA, Quek L, et al. M2 macrophages drive leukemic transformation by imposing resistance to phagocytosis and improving mitochondrial metabolism. Science Adv. AAAS; 2023;9.

58. Rutella S, Vadakekolathu J, Maziotta F, Reeder S, Yau TO, Mukhopadhyay R, et al. Immune dysfunction signatures predict outcomes and define checkpoint blockade-unresponsive microenvironments in acute myeloid leukemia. J Clin Invest.; 2022; 132(21).

59. Pospori C, Grey W, Gonzalez Anton S, Gibson S, Georgiou C, Birch F, et al. Dynamic regulation of hierarchical heterogeneity in Acute Myeloid Leukemia serves as a tumor immunoevasion mechanism. BioRxiv 2020.

