## Supplementary Figures and Tables for "SIRPα+ PD-L1+ bone marrow macrophages aid AML growth by modulating T cell function"

Supp. Figure 1: Morphology of Macrophage with AML

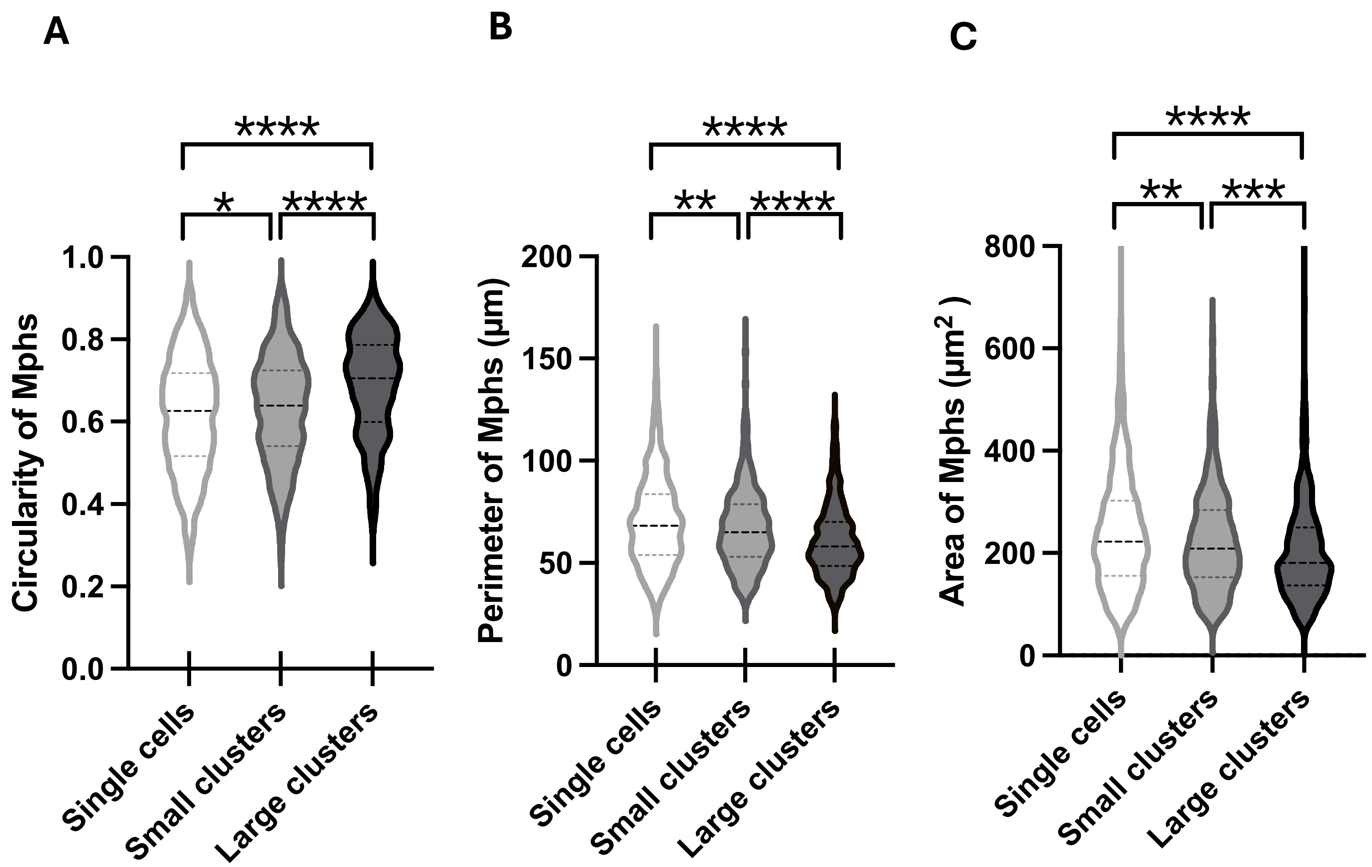

### Supp. Figure 2: Macrophage populations have different dynamics with AML

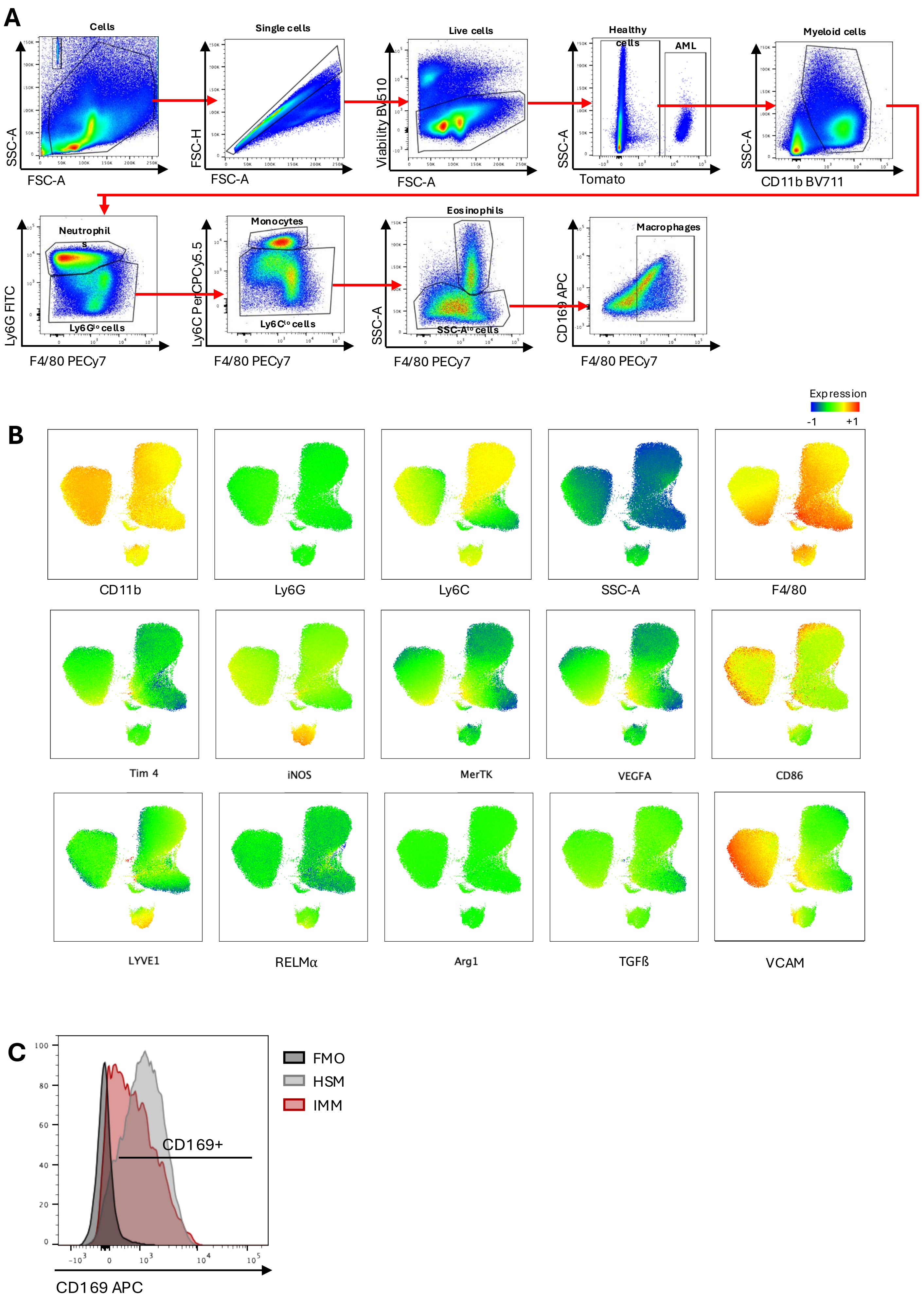

**A**

| Sample | Number of cells | Median genes per cell |
|---|---|---|
| Control macrophages | 9856 | 2574 |
| AML macrophages | 10389 | 2499 |

**B**

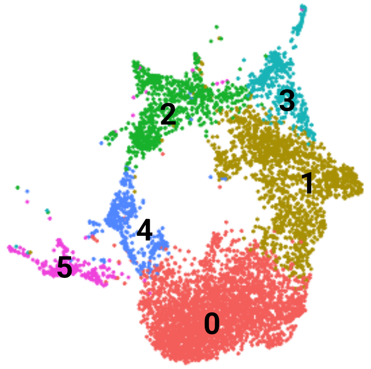


**C**

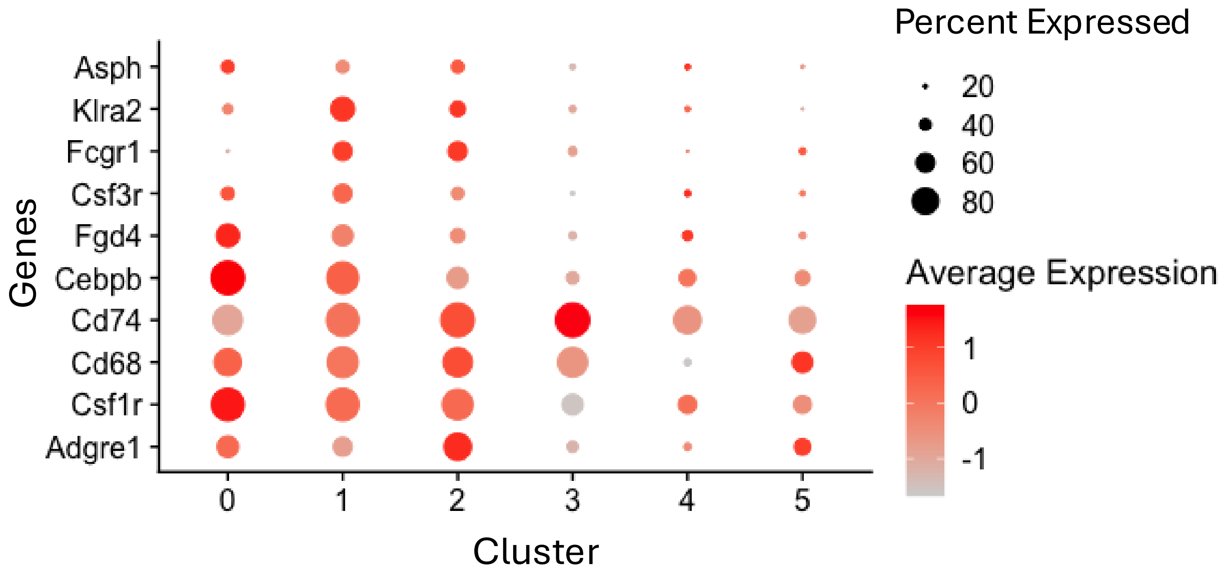


**D**

**E**

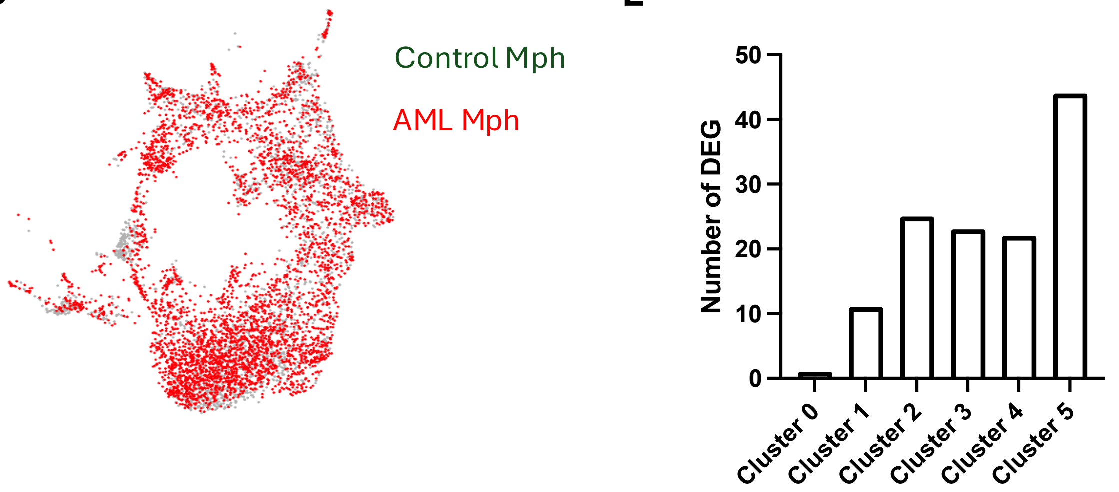

### Sup. Figure 4: Macrophage efferocytose AML cells in vivo

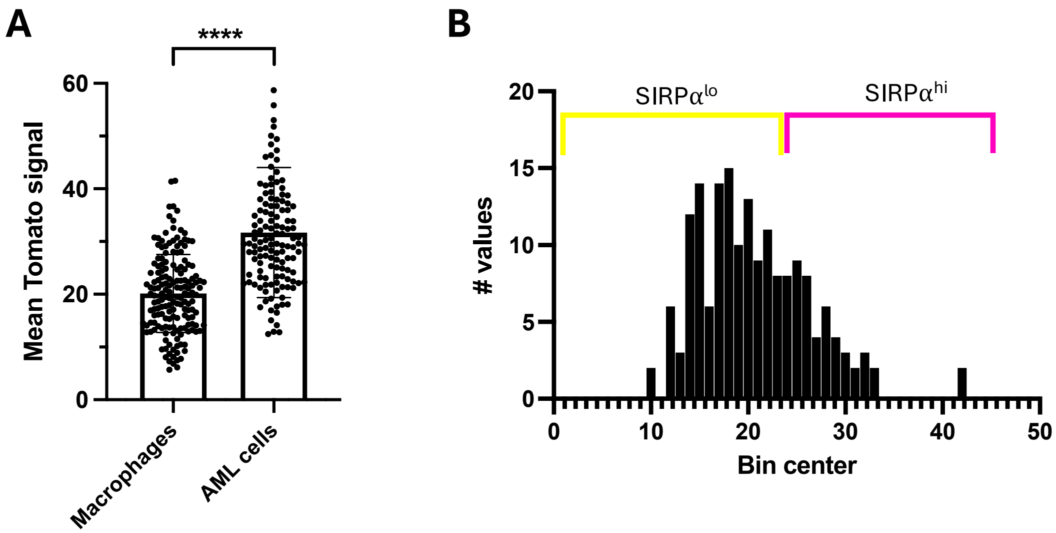

Sup. Figure 5: Ly6C$^{hi}$ monocytes differentiate towards IMMs and are capable of efferocytosing AML cells *in vitro*

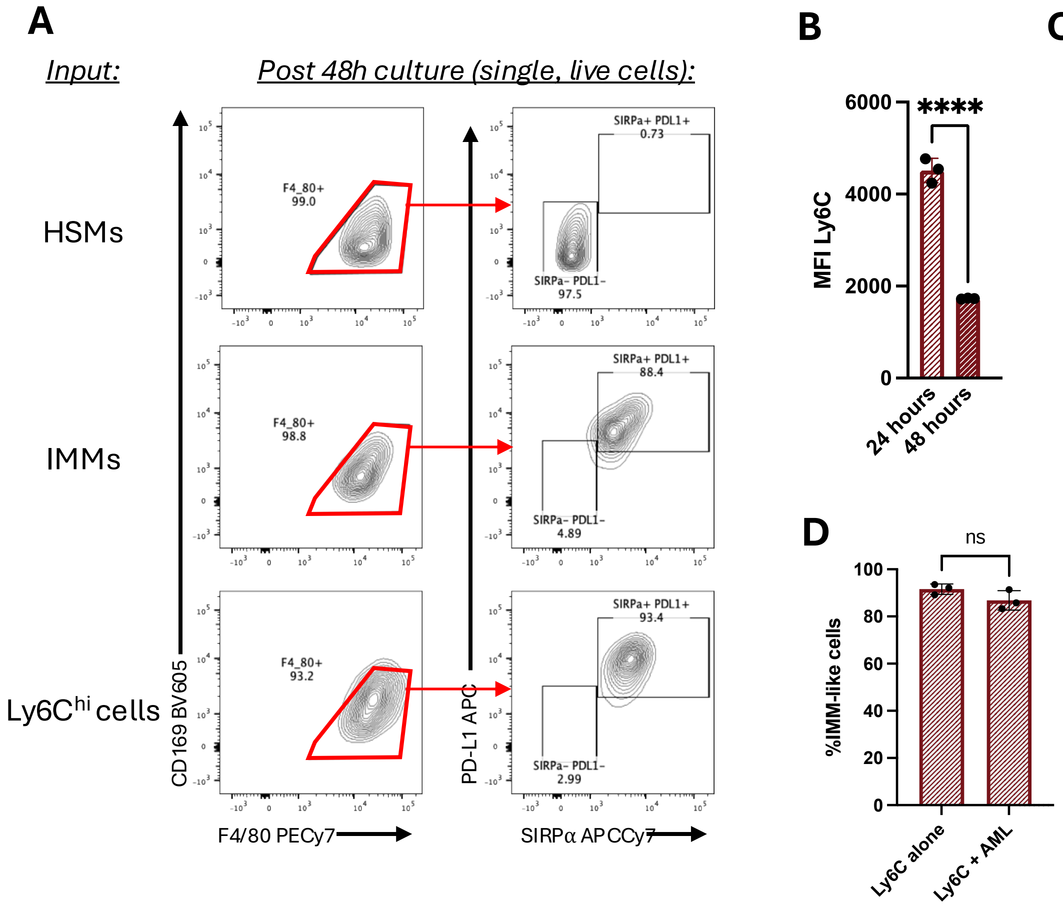

Sup. Figure 6: IMMs support AML growth by inhibiting T cells

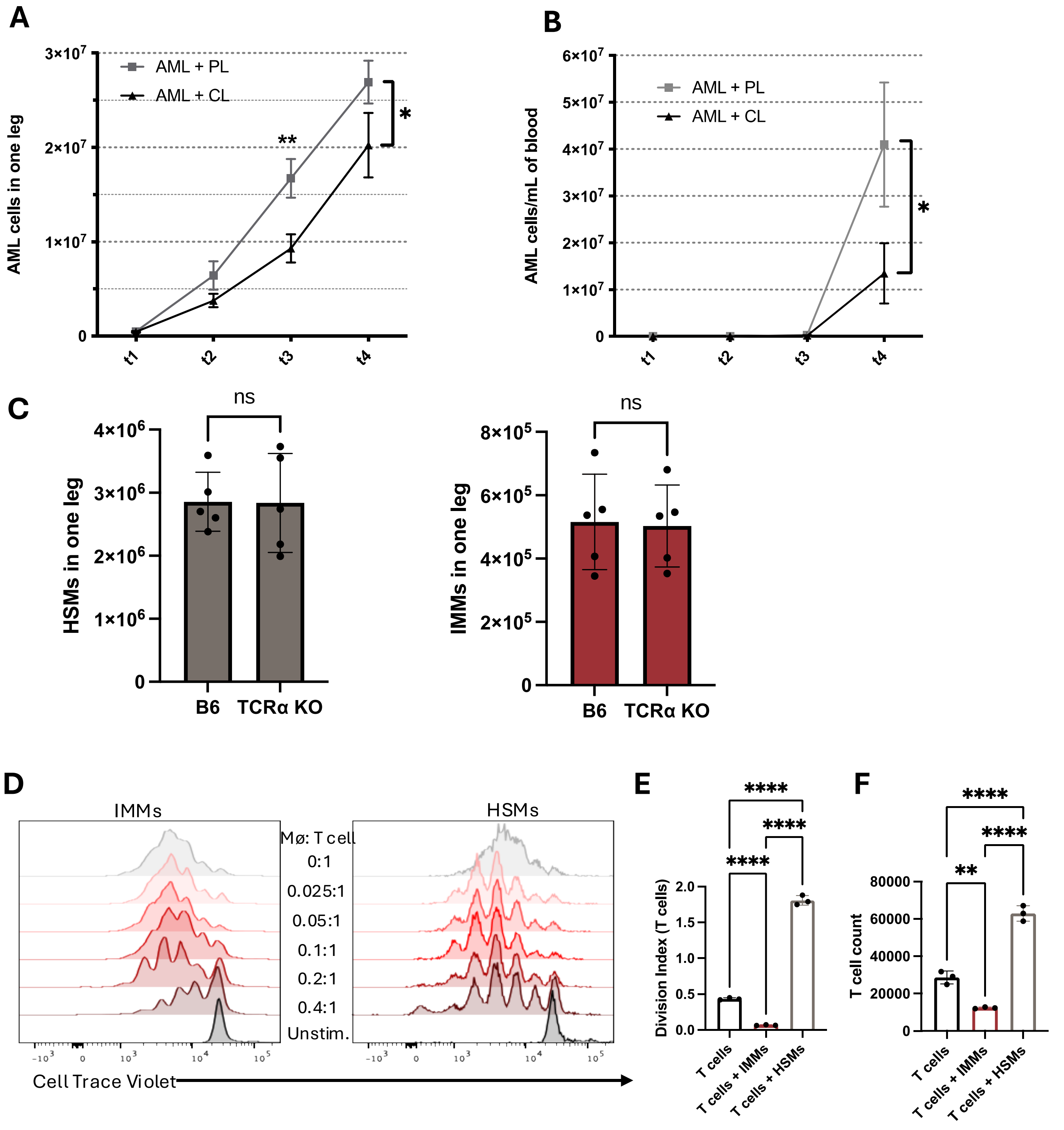

### Sup. Figure 7: IMMs are enriched post-chemotherapy

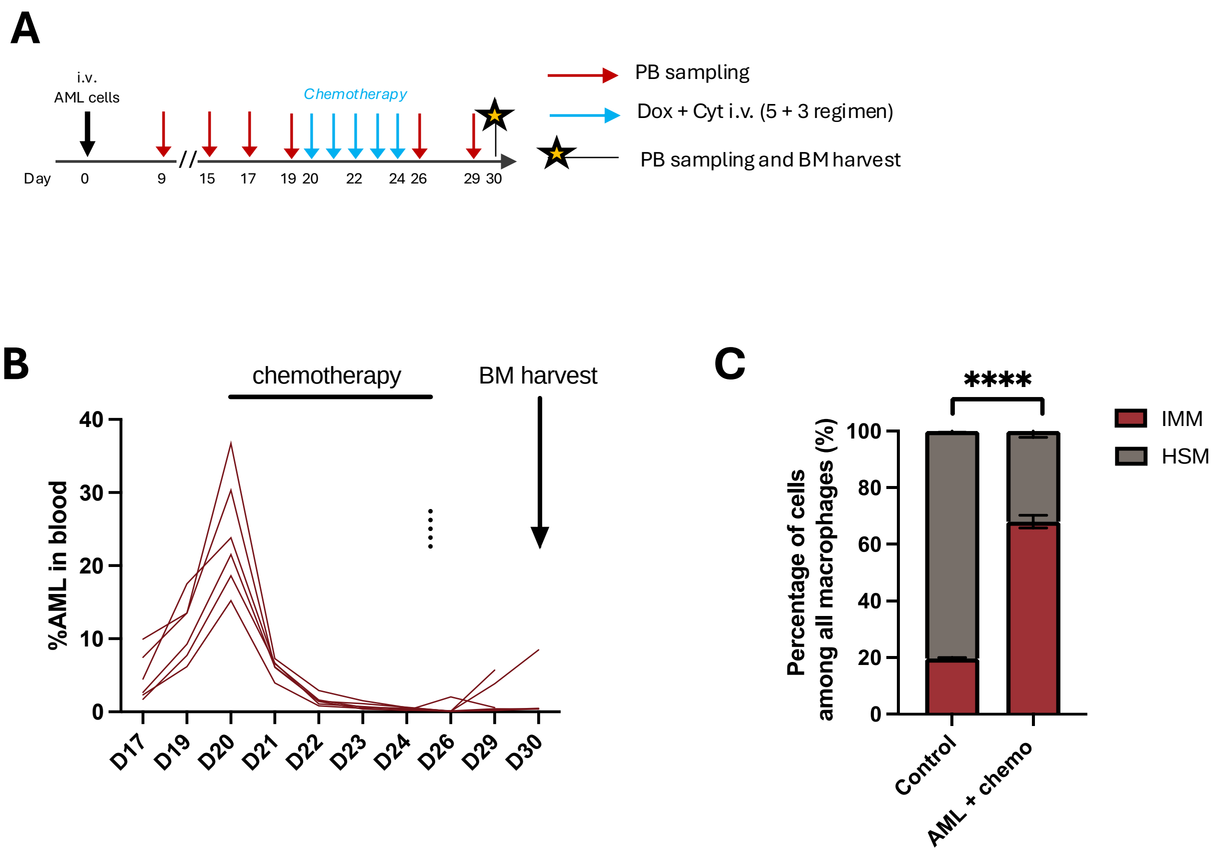

### Sup. Figure 8: Human CD206+ macrophages are similar to murine IMMs

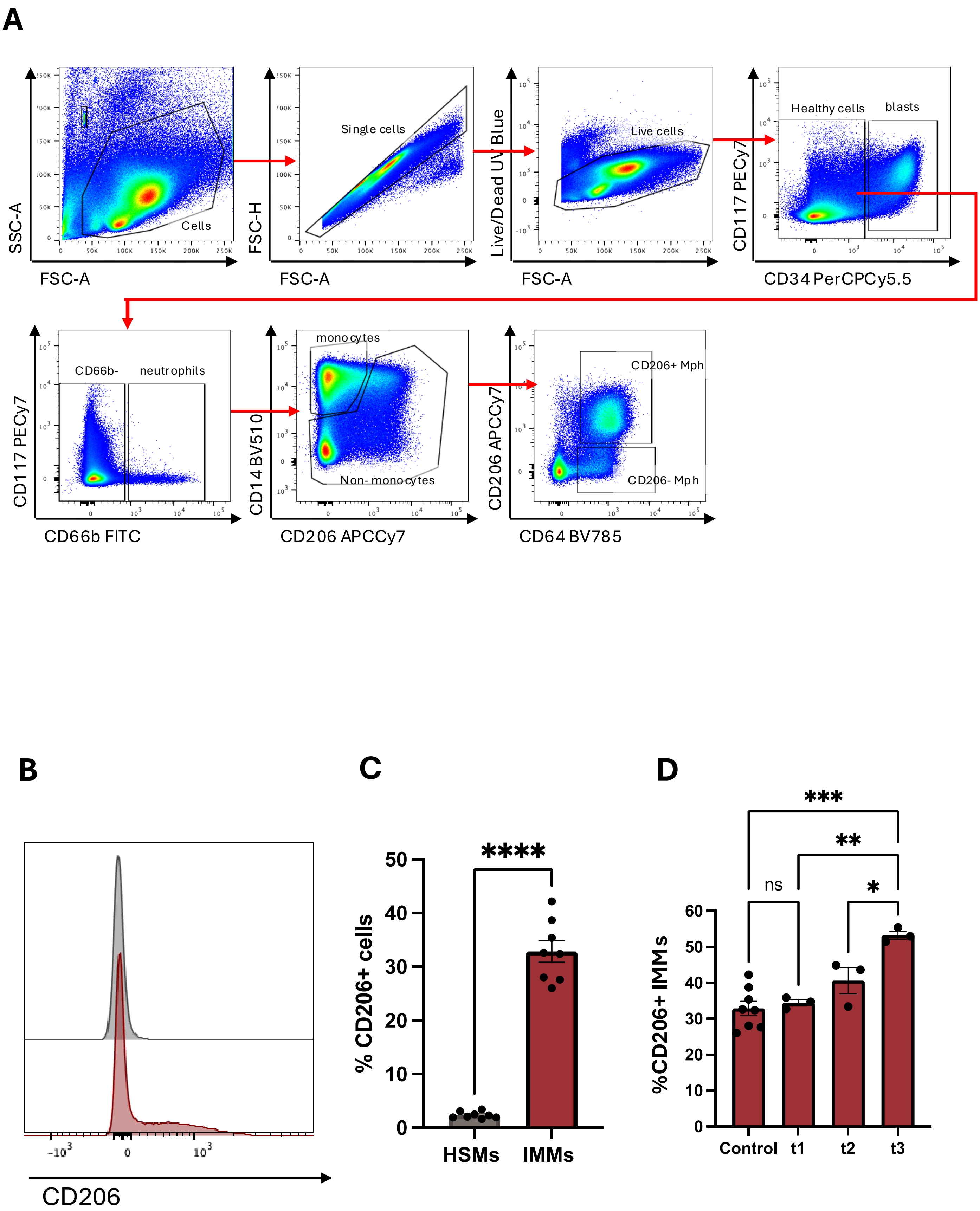

#### Sup. Table S1: Murine antibodies

| Antibody | Clone | Dilution | Source | Cat. No. |
|---|---|---|---|---|
| BV711 anti-mouse CD11b | M1/70 | 1/200 | Biolegend | 101241 |
| BUV563 anti-mouse CD11b | M1/70 | 1/500 | Biolegend | 741242 |
| FITC anti-mouse Ly6G | 1A8 | 1/500 | Biolegend | 127605 |
| BUV395 anti-mouse Ly6G | 1A8 | 1/200 | BD Biosciences | 565964 |
| PerCPCy5.5 anti-mouse Ly6C | HK1.4 | 1/600 | Biolegend | 128011 |
| BV510 anti-mouse Ly6C | HK1.4 | 1/600 | Biolegend | 128033 |
| PE/Cy7 anti-mouse F4/80 | BM8 | 1/200 | Biolegend | 123114 |
| BUV805 anti-mouse F4/80 | T45-2342 | 1/100 | BD Biosciences | 749282 |
| APC anti-mouse CD169 | 3D6.112 | 1/200 | Biolegend | 142418 |
| BV605 anti-mouse CD169 | 3D6.112 | 1/200 | Biolegend | 142413 |
| APC/Cy7 anti-mouse SIRPα | P84 | 1/200 | Biolegend | 144018 |
| PerCPCy5.5 anti-mouse SIRPα | P84 | 1/100 | Biolegend | 144010 |
| BV421 anti-mouse PD-L1 | 10F.9G2 | 1/200 | Biolegend | 124315 |
| PE-Dazzle594 anti-mouse PD-L1 | 10F.9G2 | 1/200 | Biolegend | 124323 |
| AF488 anti-mouse iNOS | 54/iNOS | 1/200 | ThermoFisher | 53-5920-80 |
| AF647 anti-mouse TNFα | MP6-XT22 | 1/100 | Biolegend | 506314 |
| PerCP-e710 anti-mouse Arg1 | A1ex5 | 1/100 | ThermoFisher | 46-3697-80 |
| AF700 anti-mouse RELMα | DS8RELM | 1/50 | ThermoFisher | 56-5441-80 |
| eFluor450 anti-mouse LYVE-1 | ALY7 | 1/20 | ThermoFisher | 48-0443-80 |
| APC/Cy7 anti-mouse MerTK | 2B10C42 | 1/200 | Biolegend | 151519 |
| BV480 anti-mouse MVCAM.A | 429 | 1/100 | BD Biosciences | 746326 |
| BV711 anti-mouse Tim-4 | RMT4-54 | 1/100 | BD Biosciences | 745509 |
| BUV737 VEGF-R2 | Avas | 1/40 | BD Biosciences | 741797 |
| BUV496 anti-mouse CD86 | PO3 | 1/100 | BD Biosciences | 750437 |
| BV421 anti-mouse LAP | TW7-16B4 | 1/100 | BD Biosciences | 565638 |
| APC anti-mouse CD3 | 145-2C11 | 1/200 | Biolegend | 100311 |
| BV650 anti-mouse CD4 | RM4-5 | 1/200 | Biolegend | 100555 |
| APC/Cy7 anti-mouse CD8 | 53-6.7 | 1/200 | Biolegend | 100714 |
| FITC anti-mouse CD44 | IM7 | 1/200 | Biolegend | 103006 |
| PerCPCy5.5 anti-mouse CD62L | MEL-14 | 1/200 | Biolegend | 104431 |

### Sup. Table S2: Human antibodies

| Antibody | Clone | Dilution | Source | Cat. No. |
|---|---|---|---|---|
| FITC anti-human CD66b | G10F5 | 1/200 | Biolegend | 305103 |
| BV510 anti-human CD14 | M5E2 | 1/200 | Biolegend | 301841 |
| PerCP/Cy5.5 anti-human CD16 | 3G8 | 1/200 | Biolegend | 302027 |
| PE/Cy7 anti-human HLA-DR | LN3 | 1/200 | Biolegend | 327017 |
| APC anti-human CD169 | 7-239 | 1/200 | Biolegend | 346007 |
| BV421 anti-human CD163 | GHI/61 | 1/200 | Biolegend | 333611 |
| APC/Cy7 anti-human CD206 | 15-2 | 1/200 | Biolegend | 321119 |
| PE anti-human SIRPα | 15-414 | 1/200 | Biolegend | 372103 |
| BV650 anti-human PD-L1 | 29E.2A3 | 1/200 | Biolegend | 329739 |
| BV785 anti-human CD64 | 10.1 | 1/200 | Biolegend | 305043 |
| PE/Cy7 anti-human CD117 | 104D2 | 1/200 | Biolegend | 313211 |
| PerCP/Cy5.5 anti-human CD34 | 581 | 1/200 | Biolegend | 343521 |
| PE/Cy7 anti-human CD64 | X54-5/7.1 | 1/200 | Biolegend | 139314 |

### Sup. Table S3: Details of AML patient samples used (London).

| Sample No. | | Age at diagnosis | AML Diagnosis | Mutations at diagnosis | Cytogenetics at Diagnosis | Treatment | Disease status at time of follow-up sample | Outcome |
|---|---|---|---|---|---|---|---|---|
| **981**<br>**Good risk** | P 1 | 19 | AML inv(16)(p13;q22) | NRAS p.Gln61Arg (VAF 39%) | CBFB::MYH11 fusion | DA + GO<br>DA + GO<br>HiDAC<br>HiDAC | Complete remission<br>MRD positive | Ongoing molecular remission |
| **975**<br>**Poor risk** | P 2 | 44 | AML with mutated FLT3 | WT1 p.Ala365ValfsTer4 (VAF 47%) variant | FISH: Trisomy 8 | DA + midostaurin<br>DA + midostaurin<br>FLA-Ida | Complete remission<br>MRD positive | Relapsed |
| **947**<br>**Good risk** | P 3 | 17 | AML with mutated NPM1 | DNMT3A p.Arg771Gln (VAF 46%), NPM1 p.Trp288CysfsTer12 (VAF 25%), WT1 p.Arg352AspfsTer6 (VAF 20%), NRAS p.Gly12Asp (VAF 16%) and NRAS p.Gly13Val (VAF 8%) variants | FISH panel negative, CMA negative | DA + GO<br>DA<br>HiDAC<br>HiDAC | Complete remission<br>MRD positive | Ongoing molecular remission |
| **928**<br>**Poor risk** | P 4 | 46 | AML NOS | NRAS p.Gly13Asp (VAF 65%) and WT1 p.His390ArgfsTer43 (VAF 30%) | FISH panel negative | DA + GO | Refractory Disease | Primary Refractory Disease |
| **1096**<br>**Poor risk** | P 5 | 16 | AML NOS | PHF6 p.Arg129Ter (VAF 37%) and WT1 p.Arg463Ter (VAF 92%) variants. A KMT2A PTD is detected. | FISH panel negative, CMA negative | MYECHILD (MA+GO)<br>FLA-Ida<br>FLA<br>Bu/Cy ASCT | Complete remission<br>MRD positive | CR but persistent high-level MRD |
| **1076**<br>**Good risk** | P 6 | 50 | AML t(8;21)(q22;q22) | KIT p.Asn822Lys (VAF 17%) and NRAS p.Gln61Arg (VAF 24%) variants. A RUNX1::RUNX1T1 fusion is detected. | RUNX1::RUNX1 T1 fusion gene accompanied by trisomy 8 and loss of 1 homolog of X. | DA + GO<br>DA + GO<br>HiDAC<br>HiDAC | Complete remission<br>MRD positive | Died in remission due to treatment complications |

### Sup. Table S4: Details of AML patient samples used (Lausanne).

| Sample No. | | Age at diagnosis | AML Diagnosis | Mutations at diagnosis | Cytogenetics at Diagnosis | Treatment | Outcome |
|---|---|---|---|---|---|---|---|
| **017** **Good risk** | P7 | 59 | AML with mutated NPM1 | NPM1 mut, WT1 overexpression | 46,XY | DA, DA, EM | Complete remission |
| **100** **Poor risk** | P8 | 63 | AML NOS, FLT3-ITD pos NPM1 neg | FLT3-ITD pos | 46,XY | FLAG+Mido, ECLOA, CLAGE, AZAVEN, allo | Relapsed |
| **103** **Poor risk** | P9 | 57 | AML with MDS related changes | IHD2 mut, WT1 overexpression | complex caryotype | IA, DA, AZAVEN, allo | Complete remission |
| **134** **Good risk** | P10 | 45 | AML with CBFC-MYH11 / inv16 and FLT3-ITD pos | CBFB-MYH11 (81%), and FLT3-ITDlow (insertion 27 bp) VAF 2.9%, KIT D816V VAF 5%, KRAS G12D VAF 2.8&, Q61H VAF 13% | 46,XX,inv(16) | IA, DA, allo | Relapsed |
| **141** **Poor risk** | P11 | 68 | AML therapy related | EZH2 mut, PRPF8 mut, RUNX1 mut (2 different ones), YRSR2 mut | 46,XY | DA, A, AZA, allo | Complete remission |
| **172** **Poor risk** | P12 | 75 | AML with MDS related changes | ASXL1 mut (Q748*), KRAS mut (G12D), TET2 mut (2 different ones) | 47,XY,+13 | FLAG, FLAG, AZAVEN | Progressive disease |
| **175** **Poor risk** | P13 | 52 | AML NOS with mutations in FLT3TKD, IDH2, SRSF2 and WT1 overexpression | FLT3-TKD D835V, IDH2 R140Q, SRSF2 P95_R102del, WT1 overexpresison | 92,XXYY[6], 46,XY[16] | IA , FLAG, AZA+Mido, allo | Complete remission |
| **194** **Poor risk** | P14 | 60 | AML NOS, AML without maturation (M1 according to FAB ) | FLT3-ITD pos (high), IDH2 mut, RUNX1 mut, KMT2a partial tandem duplication | 46,XX | IA+Mido,FLAG, AZA, allo | Complete remission |
| **195** **Poor risk** | P15 | 68 | AML with myelodysplasia related changes | CEBPA mut, GATA2 mut, EZH2 loss, CEBPA LOH, and additional unspecified chromosomal losses | Complex caryotype | IA, AZA, FLAG, allo | Complete remission |
